# Inferring spatial single-cell-level interactions through interpreting cell state and niche correlations learned by self-supervised graph transformer

**DOI:** 10.1101/2024.08.21.608964

**Authors:** Xiao Xiao, Le Zhang, Hongyu Zhao, Zuoheng Wang

**Author notes:** Correspondence Zuoheng Wang.

## Abstract

Cell-cell interactions (CCI), driven by distance-dependent signaling, are important for tissue development and organ function. While imaging-based spatial transcriptomics offers unprecedented opportunities to unravel CCI at single-cell resolution, current analyses face challenges such as limited ligand-receptor pairs measured, insufficient spatial encoding, and low interpretability. We present GITIII, a lightweight, interpretable, self-supervised graph transformer-based model that conceptualizes cells as words and their surrounding cellular neighborhood as context that shapes the meaning or state of the central cell. GITIII infers CCI by examining the correlation between a cell’s state and its niche, enabling us to understand how sender cells influence the gene expression of receiver cells, visualize spatial CCI patterns, perform CCI-informed cell clustering, and construct CCI networks. Applied to four spatial transcriptomics datasets across multiple species, organs, and platforms, GITIII effectively identified and statistically interpreted CCI patterns in the brain and tumor microenvironments.

## Introduction

In multicellular organisms, cell-cell interactions (CCI) play a pivotal role in maintaining tissue development and organ function. These interactions may occur through direct physical contact mediated by cell adhesion molecules between adjacent cells or through longer-range paracrine and autocrine signaling, where secreted factors activate intracellular signaling cascades that modulate gene expression and cellular behavior. CCI are influenced by multiple factors, including the spatial proximity of interacting cells, the local tissue microenvironment of the receiver cell, the strength of ligand-receptor signaling, and the influence of gene regulatory networks on downstream responses. Together, these factors orchestrate cell function and tissue homeostasis.

Single-cell RNA sequencing (scRNA-seq) and spatial transcriptomics (ST) have offered unprecedented opportunities to unravel intercellular communication. Many existing methods focus on constructing CCI networks between cell types by identifying ligand-receptor pairs through which communication occurs^1^. These approaches span statistical tools for detecting significant interactions^2–6^, and techniques that incorporate spatial information through graph neural networks (GNN), optimal transport, or distance-based weighting and thresholding^7–18^. Collectively, they have greatly advanced our understanding of ligand-receptor-mediated molecular interactions^6,19–23^. However, most of these methods treat cells of the same type as homogeneous, often overlooking within-cell-type heterogeneity. In reality, cells of the same cell-type pair may interact only when spatially proximal, but not when distant, due to microenvironmental differences or spatial constraints. Moreover, current approaches tend to focus only on the most statistically extreme communication signals, potentially missing subtle interactions that still have downstream effects on gene expression and cell behavior.

ST, which captures both gene expression and the spatial location of cells within tissue, has become a powerful tool for studying CCI at single-cell resolution, particularly in the context of the tissue microenvironment. However, identifying CCI from imaging-based ST^24–26^ remains challenging, as these platforms typically measure only a limited set of genes and lack adequate coverage of ligand-receptor pairs. Thus, many CCI analysis tools that rely on ligand-receptor information to detect significant interactions are not applicable to such data. Additionally, spatial information has not been fully exploit in current CCI analysis methods. Some approaches either restrict CCI inference to a small predefined spatial neighborhood^7,14,27,28^, or fail to account for the effect of distance on interaction strength^8^, or use a fixed distance-scaling model across all signaling pathways^10,15,29^, despite the fact that different ligand-receptor signaling may exhibit distinct distance-dependent patterns in biological systems. Furthermore, GNN-based methods for CCI analysis^30–32^ often suffer from inadequate graph encoding and limited interpretability, particularly when using multi-layer architecture. These limitations hinder biological interpretation and reduce their utility for uncovering mechanistic insights into CCI.

An important aspect of CCI inference is to estimate the effects of neighboring sender cells on the gene expression of receiver cells through CCI that, to our knowledge, has not been explicitly studied in previous studies. We introduce GITIII (Graph Inductive bias Transformer for Intercellular Interaction Investigation), an interpretable, self-supervised graph transformer-based language model that treats cells as words and their surrounding cellular neighborhood as a sentence to explore CCI within a cellular network using their spatial and transcriptomic information from imaging-based ST. GITIII infers spatial single-cell-level interactions by examining the correlation between a cell’s state and the spatial organization, ligand expression, and the types and states of its neighboring cells. Its interpretable architecture allows us to estimate the impact of sender cells on the gene expression of receiver cells, visualize spatial patterns of CCI, perform CCI-informed cell clustering, construct CCI networks, and compare CCI strength across conditions. Applied to four diverse ST datasets, GITIII successfully identified and quantitatively interpreted CCI patterns that drive within-cell-type heterogeneity, thus advancing our understanding of brain structure and the tumor microenvironment (TME).

## Results

### Overview of GITIII

GITIII is a lightweight and interpretable deep learning model for inferring spatial single-cell-level interaction patterns. It takes gene expression, spatial coordinates, and cell type annotations as inputs. The model first decomposes the gene expression levels of each cell into two parts: “cell-type expression,” representing the average gene expression across cells of the same type, and “cell-state expression,” representing the deviation of its gene expression from the average expression of its cell type (Fig. 1a). For each cell, GITIII constructs a cellular neighborhood of size 𝑘 based on spatial proximity and learns the interactions between the central cell and its neighboring cells. The cellular neighborhood is represented as a spatial subgraph, where nodes are cells and edges are spatial proximity between cells, or alternatively, as a sentence where cells are words and neighboring cells provide context to shape the meaning or state of the central cell. GITIII employs a single-layer graph transformer to learn interaction patterns and computes an influence tensor that quantifies the impact of neighboring cells on the cell-state expression of the central cell (Fig. 1b, 1c).

**Fig. 1.**
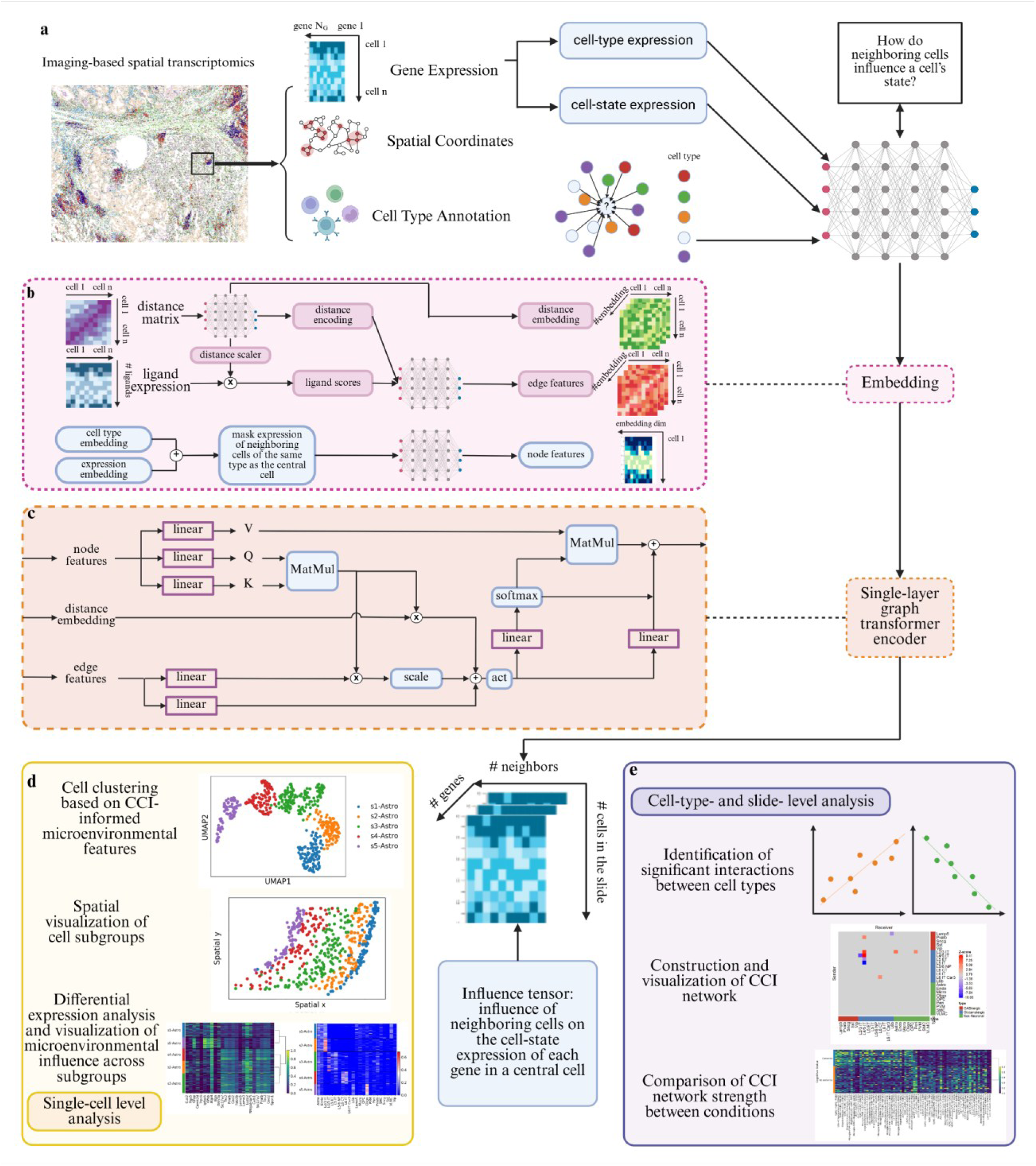
Schematic diagram of GITIII. **a,** GITIII Workflow. The model employs self-supervised learning to infer cell-cell interactions (CCI) by predicting the cell-state expression of the central cell from the spatial organization, ligand expression, and the gene expressions of its neighboring cells. We then estimate the influence of neighboring cells on the cell-state expression of the central cell, and perform downstream analyses. **b,** Detailed view of the input data embeddings. Node features incorporate cell type embeddings and expression encodings, and edge features contain ligand gene expression scaled by distance and the embedding of distance between cells. **c,** Detailed view of the encoder layer. A graph attention mechanism designed to better capture the CCI patterns in the cellular neighborhood. Edge features are updated through linear transformations combined with the attention matrix from the node features and the embedding of distance relationships. Node features are aggregated through attention mechanisms, augmented by edge features weighted by attention scores. **d,** Single-cell level analysis. Clustering cells into subgroups based on CCI-informed microenvironmental features and spatial visualization. Differential expression analysis to interpret differences between subgroups. **e,** Cell-type level analysis. CCI network analysis to identify significant interactions between cell types. Comparison of CCI network strength between conditions.

To investigate CCI patterns across spatial regions, the influence tensor is used to visualize single-cell-level spatial patterns of CCI by plotting the spatial distance and strength of influence between interacting cells. We then perform cell clustering within each cell type using CCI-informed microenvironmental features (Fig. 1d). At the cell-type level, significant interactions between cell types are identified using Pearson correlation analysis. For each gene, CCI networks are constructed by identifying sender cell types that significantly influence the expression level of this gene in receiver cell types. At the sample level, we compare CCI network strength between conditions using a microenvironment-adjusted network strength metric (Fig. 1e).

### GITIII reveals spatial patterns of CCI in mouse brains

We illustrated the main features of GITIII using a mouse primary motor cortex (PMC) ST dataset generated by MERFISH^33^. Slice 201 from mouse 1 was used for demonstration (Fig. 2a). As expected from physical principles, the locally estimated scatterplot smoothing (LOESS) curve showed that the estimated CCI strength decreases with increasing distance between interacting cells (Fig. 2b).

**Fig. 2.**
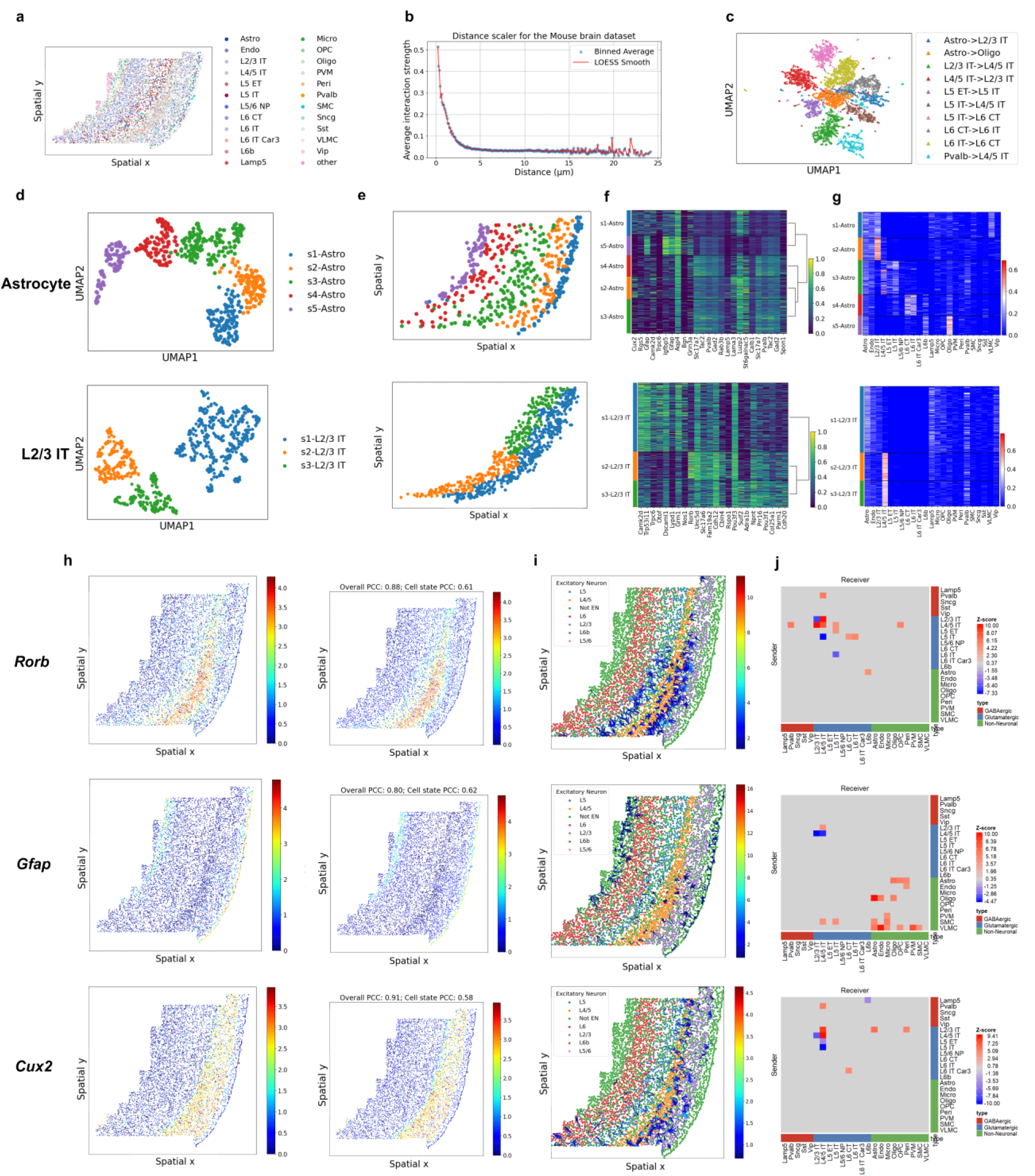
Spatial patterns of CCI in the mouse PMC dataset. **a,** Spatial distribution of 24 cell types. **b,** The LOESS curve of the estimated CCI strength decreased with increasing distance between interacting cells. **c,** UMAP visualization of CCI pairs. **d,** UMAP visualization of astrocyte (upper) and L2/3 IT neuron (lower) subgroups. **e,** Spatial distribution of astrocyte (upper) and L2/3 IT neuron (lower) subgroups. **f,** Heatmap of the top differentially expressed genes in astrocyte (upper) and L2/3 IT neuron (lower) subgroups. **g,** Neighboring cell type influence on astrocyte (upper) and L2/3 IT neuron (lower) subgroups. **h,** Visualization of the observed (left) and predicted (right) expression levels of *Rorb*, *Gfap*, and *Cux2*. **i,** Spatial distribution of CCI, showing the top 5% strongest CCI pairs influencing the expression of *Rorb*, *Gfap*, and *Cux2*, with arrows connecting the corresponding sender and receiver cells, and color representing the influence value of the interaction. In the top panel, many CCI pairs between L4/5 IT and L2/3 IT cells, particularly with L2/3 IT subtype 2, as well as CCI pairs within layer 5, influence the expression of *Rorb*. In the middle panel, many CCI pairs, involving non-neuronal cells located in layer 1 and the white matter, influence the expression of *Gfab*. In the bottom panel, CCI pairs, located in layers 2 and 3, influence the expression of *Cux2*. **j,** CCI networks influencing the expression of *Rorb*, *Gfap*, and *Cux2*, with color representing the z-score of the Pearson correlation of interactions between cell types that influence the gene expression of a receiver cell type, positive values indicating upregulation and negative values indicating downregulation.

Single-cell-level interactions within the same cell-type pair are expected to be more similar than interactions across different pairs, because the ligand-receptor expression patterns in sender-receiver pairs, as well as the downstream gene regulatory responses in receiver cells, are generally more similar within a given cell-type combination. Fig. 2c shows that the 10 most frequent cell-type interactions from the selected CCI pairs across all cellular neighborhoods formed distinct clusters in the Uniform Manifold Approximation and Projection (UMAP), where each point represents one CCI pair. This confirms that CCI pairs from the same cell-type combination have similar patterns and that GITIII effectively distinguishes interactions among different cell types.

We then used CCI-informed microenvironmental features to cluster astrocytes and L2/3 IT neurons. Astrocytes are known to display layer-specific morphological and molecular differences^34^ and to be functionally diverse across brain regions^35^. In the MERFISH study^33^, astrocytes were clustered into three subgroups: one distributed across all layers and two enriched mostly in the white matter (Extended Data Fig. 1b). In comparison, GITIII identified five subgroups with clear layer-specific distributions (Fig. 2d), where subgroup 1 located in layer 1, subgroup 2 located in layers 2 and 3, and subgroup 5 located in the white matter (Fig. 2e, Extended Data Fig. 1a). This pattern is consistent with the reported heterogeneity of astrocytes across cortical layers in both mouse and human brains^36–39^. Subgroups 1 and 2 showed increased expression of *Cux2* and *Lama3*, while subgroup 5 showed increased expression of *Igfbp5* and *Gfap* (adjusted 𝑝 < 1 × 10^−5^, Fig. 2f, Extended Data Fig. 1d), suggesting that GITIII captured microenvironment-driven transcriptional variation. We also observed *Luzp2* expression in several subgroups, consistent with a thalamus-enriched (*LUZP2*^+^*DCLK1*^+^) astrocyte subtype found in the human brain^40^. GITIII further revealed that subgroups 1 and 2 were strongly influenced by L2/3 IT neurons, and subgroup 5 was primarily influenced by oligodendrocytes and L6b neurons (Fig. 2g), confirming that intercellular communication is most likely to occur between cells in close physical proximity (Extended Data Fig. 1a, 1c).

For L2/3 IT neurons, the MERFISH study^33^ identified five subgroups without clear spatial boundaries (Extended Data Fig. 1f). In contrast, GITIII detected three subgroups with clear separations (Fig. 2d). Subgroup 1 located in the upper region of layers 2 and 3, closer to layer 1, while subgroups 2 and 3 located in the deeper region of layers 2 and 3, closer to layer 4 (Fig. 2e, Extended Data Fig. 1g). Subgroup 1 showed upregulated expression of *Camk2d*, *Trp53i11*, *Trpc6*, and *Otof*, subgroup 2 showed increased expression of *Rorb* and *Unc5d*, while subgroup 3 showed increased expression of *Sulf2* and *Adra1b* (adjusted 𝑝 < 1 × 10^−2^^4^, Fig. 2f, Extended Data Fig. 1h). GITIII inferred that subgroup 1 was primarily influenced by astrocytes, Lamp5, Pvalb, and Vip neurons, while subgroups 2 and 3 were strongly influenced by L4/5 IT neurons as they were physically closer to layer 4, and subgroup 3 was also influenced by Pvalb neurons (Fig. 2g).

To compare our subgroup results with spatial clustering, we applied BANKSY^41^, which clusters all cells based on their transcriptomes and local microenvironment. Like other domain detection methods, BANKSY clusters all cells on a tissue slide, thus each cluster may include multiple cell types. In contrast, GITIII clusters within a cell type to identify subgroups that differ in their surrounding cellular neighborhood. Supplementary Fig. 2 shows the clustering results from BANKSY at different resolutions, restricted to clusters containing at least 20 astrocytes or L2/3 IT neurons. Astrocytes and L2/3 IT neurons not included in these clusters were grouped together for visualization. BANKSY detected several tissue domains containing astrocytes and L2/3 IT neurons distributed across cortical layers, especially at higher resolutions. However, GITIII showed clearer layer-specific separation among astrocyte subgroups (Fig. 2e), while BANKSY grouped a small number of astrocytes and L2/3 IT neurons with other cell types. Together, the consistent spatial separation observed in both methods supports that the subgroups identified by GITIII are driven by CCI-informed microenvironmental features.

We further visualized the inferred CCI and constructed CCI networks that influence the expression of three biologically relevant genes, *Rorb*, *Gfap*, and *Cux2*, chosen based on their high predictive accuracy. Pearson correlation coefficients between the observed and predicted cell-state expression of these genes across all cells were greater than 0.58 (Fig. 2h). We then visualized the spatial distribution of the top 5% of CCI pairs that strongly influence the expression levels in receiver cells (Fig. 2i). For example, interactions between L4/5 IT and L2/3 IT cells were associated with increased expression of *Rorb* (Fig. 2j), where we observed more CCI pairs between L4/5 IT and L2/3 IT subgroup 2 (Fig. 2i, Extended Data Fig. 1g, 1i), consistent with the upregulated expression of *Rorb* in L2/3 IT subgroup 2 (Fig. 2f). Interactions between oligodendrocytes and astrocytes were associated with increased expression of *Gfap* (Fig. 2i, 2j, Extended Data Fig. 1j), consistent with the inferred interactions between oligodendrocytes and astrocyte subgroup 5 in the white matter (Extended Data Fig. 1c) and the upregulated expression of *Gfap* in astrocyte subgroup 5 (Fig. 2f), and supporting the role of astrocyte-oligodendrocyte interaction in myelin regeneration^42^. *Cux2*, a gene encoding a transcription factor involved in neuronal proliferation and differentiation^43^, showed increased expression in astrocytes influenced by L2/3 IT neurons (Fig. 2i, 2j). This result agreed with the upregulated expression of *Cux2* in astrocyte subgroup 2, located in layers 2 and 3 (Extended Data Fig. 1a, 1k), which may related to the influence of neurons on astrocyte morphology and proliferation^44^.

### GITIII identifies CCI patterns within and across patients with Alzheimer’s disease

We analyzed the SEA-AD middle temporal gyrus ST dataset generated by MERFISH^45^. Sample ‘H20.33.001.CX28.MTG.02.007.1.02.03’ was used to demonstrate within-sample performance (Fig. 3a). As expected, interaction strength generally decreased with increasing distance between interacting cells (Fig. 3b). However, adjacent cells can have weaker interactions than more distant cells. The strength of interactions influencing the expression of *RORB* varied substantially within 18 µm and became consistently weak beyond this distance (Extended Data Fig. 2a). This pattern suggests that spatial proximity alone does not determine interaction strength. Even when two cells are physically close, low ligand-receptor expression can lead to weak interactions. The UMAP of the 10 most frequent cell-type interactions showed well-separated CCI pairs, indicating that GITIII effectively identified distinct patterns across interactions among different cell types (Fig. 3c).

**Fig. 3.**
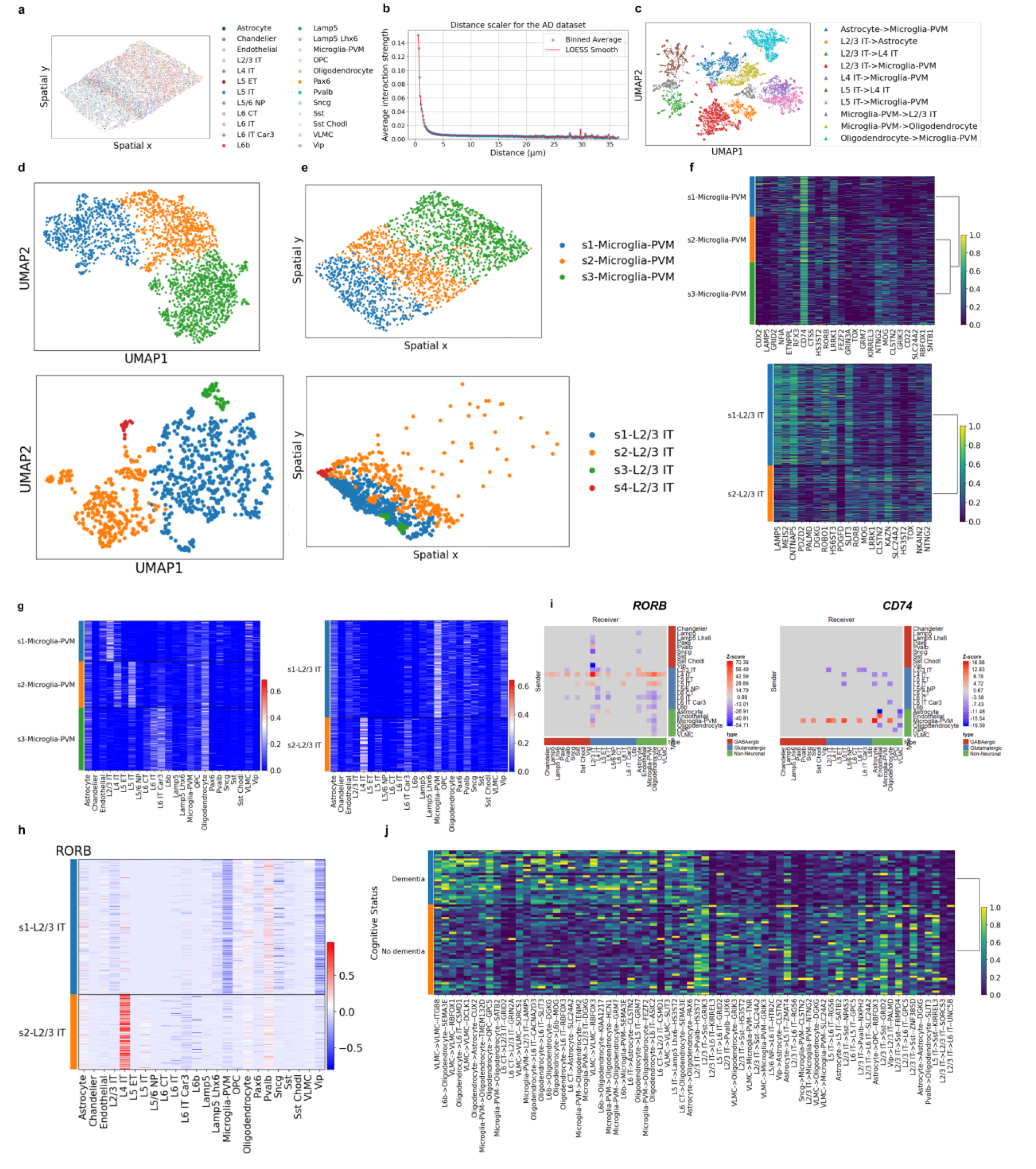
CCI patterns within and across patients in the SEA-AD dataset. **a,** Spatial distribution of 24 cell types. **b,** The LOESS curve of the estimated CCI strength decreased with increasing distance between interacting cells. **c,** UMAP visualization of CCI pairs. **d,** UMAP visualization of microglia (upper) and L2/3 IT neuron (lower) subgroups. **e,** Spatial distribution of microglia (upper) and L2/3 IT neuron (lower) subgroups. **f,** Heatmap of the top differentially expressed genes in microglia (upper) and L2/3 IT neuron (lower) subgroups. **g,** Neighboring cell type influence on microglia (left) and L2/3 IT (right) subgroups. **h,** Neighboring cell type influence on the expression of *RORB* in L2/3 IT subgroups, with positive values indicating upregulation and negative values indicating downregulation. **i,** CCI networks influencing the expression of *RORB* and *CD74*, with color representing the z-score of the Pearson correlation of interactions between cell types that influence the gene expression of a receiver cell type, positive values indicating upregulation and negative values indicating downregulation. **j,** Heatmap of the top differential CCI network strength between patients with or without dementia.

To investigate how a cell’s local microenvironment influences its state, we focused on microglia, for their involvement in neuroinflammation in AD^46,47^, and L2/3 IT neurons, a vulnerable neuronal population in AD^48^. Microglial state is correlated with laminar location in the mouse cortex^49^. The UMAP visualization of microglia using CCI-informed microenvironmental features revealed three clusters, each localized within different cortical layers (Fig. 3d, 3e, Extended Data Fig. 2b), consistent with the reported laminar organization of microglia in human ST studies^50,51^. We observed upregulated expression of *CUX2* in subgroup 1 and *NTNG2* in subgroup 3 (adjusted 𝑝 < 1 × 10^−2^^4^, Fig. 3f). GITIII further inferred that subgroup 1 was strongly influenced by L2/3 IT neurons, subgroup 2 by L4 IT and L5 IT neurons, and subgroup 3 by L6 IT and L6 IT Car3 neurons (Fig. 3g). These interactions support that intercellular communication primarily occurs between cells that are physically close (Extended Data Fig. 2b). These identified genes, whose expression was influenced by CCI, may impact microglial functions including proliferation, synapse formation^52^, accumulation along axons^53,54^, and myelin growth and integrity^55^, confirming the role of neurons in determining microglial state^56^.

GITIII clustered L2/3 IT neurons into four subgroups, with most cells belonging to the first two subgroups (Fig. 3d). Subgroups 1, 3, and 4 located in the upper region of layers 2 and 3, where most of the vulnerable neuronal populations locate^45^, whereas subgroup 2 located in the deeper region of layers 2 and 3, close to layer 4 (Fig. 3e). These subgroups had much clearer spatial separation compared to the gene expression-based subgroups (Extended Data Fig. 2c)^45^. Subgroup 1 showed downregulated expression of *RORB* (adjusted 𝑝 < 1 × 10^−2^^4^, Fig. 3f), a marker for selectively vulnerable neurons^48^, and was mostly influenced by microglia, while subgroup 2 was primarily influenced by L4 IT neurons (Fig. 3g).

We then constructed CCI networks for two AD risk genes *RORB*^48^ and *CD74*^57^ (Fig. 3i). For *RORB*, its expression in L2/3 IT subgroup 1 was downregulated by interactions with microglia and VIP neurons, and upregulated in subgroup 2 by interactions with L4 IT neurons (Fig. 3h). *CD74*, a microglia maker involved in immune responses^58^, was inferred to be significantly upregulated in cell types interacting with microglia, highlighting the influence of inflammatory microglia on neighboring cells^59^.

Finally, we quantified the CCI network strength within each slide and observed some separation among the 69 slides from patients with and without dementia (Extended Data Fig. 2d). Dementia patients had stronger interactions between microglia, oligodendrocytes, and excitatory neurons in deeper cortical layers compared to patients without dementia (adjusted 𝑝 < 0.1, Fig. 3j), indicating more pronounced microglia and oligodendrocyte dysfunction in dementia that may contribute to neuronal impairment and cognitive decline.

### GITIII interprets tumor microenvironment in NSCLC

To understand the TME and its impact on cell states, we analyzed the human non-small cell lung cancer (NSCLC) ST dataset generated by CosMx^25^. Sample ‘Lung 6’ was used for demonstration (Fig. 4a). We observed an overall trend of decreasing interaction strength with increasing distance between interacting cells (Fig. 4b). The CCI pairs among the 10 most frequent cell-type interactions showed distinct separation in a UMAP (Fig. 4c).

**Fig. 4.**
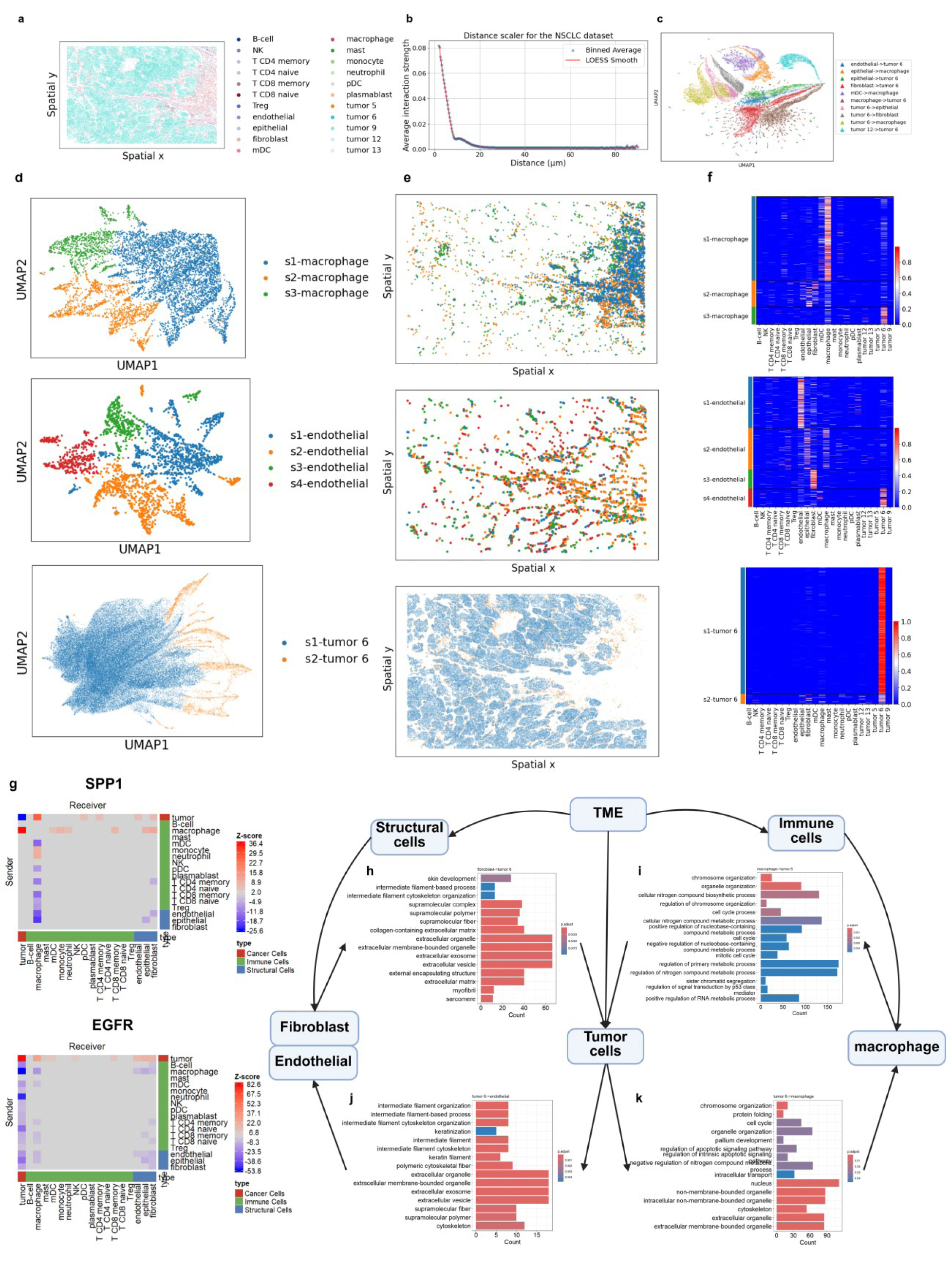
Quantitative analysis of tumor microenvironment in the NSCLC dataset. **a,** Spatial distribution of 18 cell types. **b,** The LOESS curve of the estimated CCI strength decreased with increasing distance between interacting cells. **c,** UMAP visualization of CCI pairs. **d,** UMAP visualization of macrophage (upper), endothelial cell (middle), and tumor cell (lower) subgroups. **e,** Spatial distribution of macrophages (upper), endothelial cells (middle), and tumor cells (lower) subgroups. **f,** Neighboring cell type influence on macrophages (upper), endothelial cells (middle), and tumor cells (lower) subgroups. **g,** CCI networks influencing the expression of *SPP1* and *EGFR*, with color representing the z-score of the Pearson correlation of interactions between cell types that influence the gene expression of a receiver cell type, positive values indicating upregulation and negative values indicating downregulation. **h-k,** GO enrichment analysis of genes whose expression are significantly influenced by CCI **(h)** from fibroblasts to tumor cells, **(i)** from macrophages to tumor cells, **(j)** from tumor cells to endothelial cells, and **(k)** from tumor cells to macrophages.

Using CCI-informed features, we clustered cells that play important roles in the TME, including macrophages, endothelial cells, and tumor cells. GITIII identified macrophage subgroup 3 and endothelial subgroup 4 as strongly influenced by tumor cells (Fig. 4d-4f). Macrophage subgroup 3 showed increased expression of *SPP1* (adjusted 𝑝 < 1 × 10^−2^^4^, Supplementary Fig. 3a), suggesting them as tumor-associated macrophages that contribute to tumor progression and the TME remodeling^25,60,61^. High expression of *EGFR* and several KRT genes was also observed in this subgroup. While these genes are not marker genes of macrophage, one possible explanation is that macrophages phagocytose nearby tumor cells that express *EGFR* and the KRT genes. The presence of these transcripts within macrophages could therefore reflect RNA from tumor cells. Endothelial subgroup 4 showed upregulated expression of *KRT6A*, *KRT17*, *EGFR*, and *KRT5* (adjusted 𝑝 < 1 × 10^−4^, Supplementary Fig. 3a), suggesting tumor-induced activation via EGFR signaling to promote proliferation and angiogenesis^62^. Tumor cells were clustered into two subgroups (Fig. 4d), where subgroup 2, located near the tumor boundary (Fig. 4e), interacted with macrophages and structural cells (Fig. 4f) and showed reduced expression of *GPNMB*, *EGFR*, *KRT19*, *CD9*, and *KRT15* (adjusted 𝑝 < 1 × 10^−2^^4^, Supplementary Fig. 3a). These genes are associated with cancer proliferation^63^, immunosuppression^64^, and metastasis^65^, highlighting the suppressive effect of immune cells at the tumor boundary.

We then constructed CCI networks for *SPP1* and *EGFR*. Interactions between tumor cells and macrophages were associated with increased expression of *SPP1* (Fig. 4g), consistent with the spatial distribution of macrophage subgroup 3 around tumor cells (Fig. 4e). Interactions between tumor cells and neighboring cell types, such as tumor cells, macrophages, and structural cells, were associated with increased expression of *EGFR* in these cell types, promoting tumor proliferation and metastasis^66^.

To further characterize the CCI-related biological functions and processes, we conducted Gene Ontology (GO) enrichment analysis on genes significantly influenced by interactions between tumor cells and macrophages, fibroblasts, and endothelial cells, using ClusterProfiler^67^. Fibroblasts mainly influenced tumor cells by remodeling the extracellular matrix^68^ (Fig. 4h), while tumor cells influenced endothelial cells by promoting cytoskeleton organization and fiber growth^69^ (Fig. 4j).

These interactions collectively enhanced tumor proliferation. Macrophages modulated tumor cells through pathways related to cell cycle regulation^70^, chromosome organization, and the p53 signal pathway^71^ (Fig. 4i). In contrast, tumor cells enhanced the role of macrophages in apoptotic processes^72^ and influenced their extracellular organelles^73^ (Fig. 4k). Together, these interactions revealed complex interactions in the TME and suggest potential therapeutic targets^74^.

We also performed enrichment analyses on cell-type-specific genes for tumor cells, macrophages, fibroblasts, and endothelial cells. Compared with pathways enriched in CCI-influenced genes, the pathways enriched in sender cell-type-specific genes were largely distinct (Supplementary Note 6, Supplementary Fig. 3b), suggesting that the CCI-related biological processes we found are unlikely to be artefacts of lateral spillover but instead reflect intercellular communication.

### GITIII uncovers CCI-driven within-cell-type heterogeneity in breast cancer

We analyzed a human breast cancer ST dataset generated by Xenium^26^ (Fig. 5a). The first replicate slide from sample 1 was used for demonstration. As expected, interaction strength generally decreased with increasing distance between interacting cells (Fig. 5b). The UMAP showed distinct separation among the CCI pairs from the 10 most frequent cell-type interactions (Fig. 5c).

**Fig. 5.**
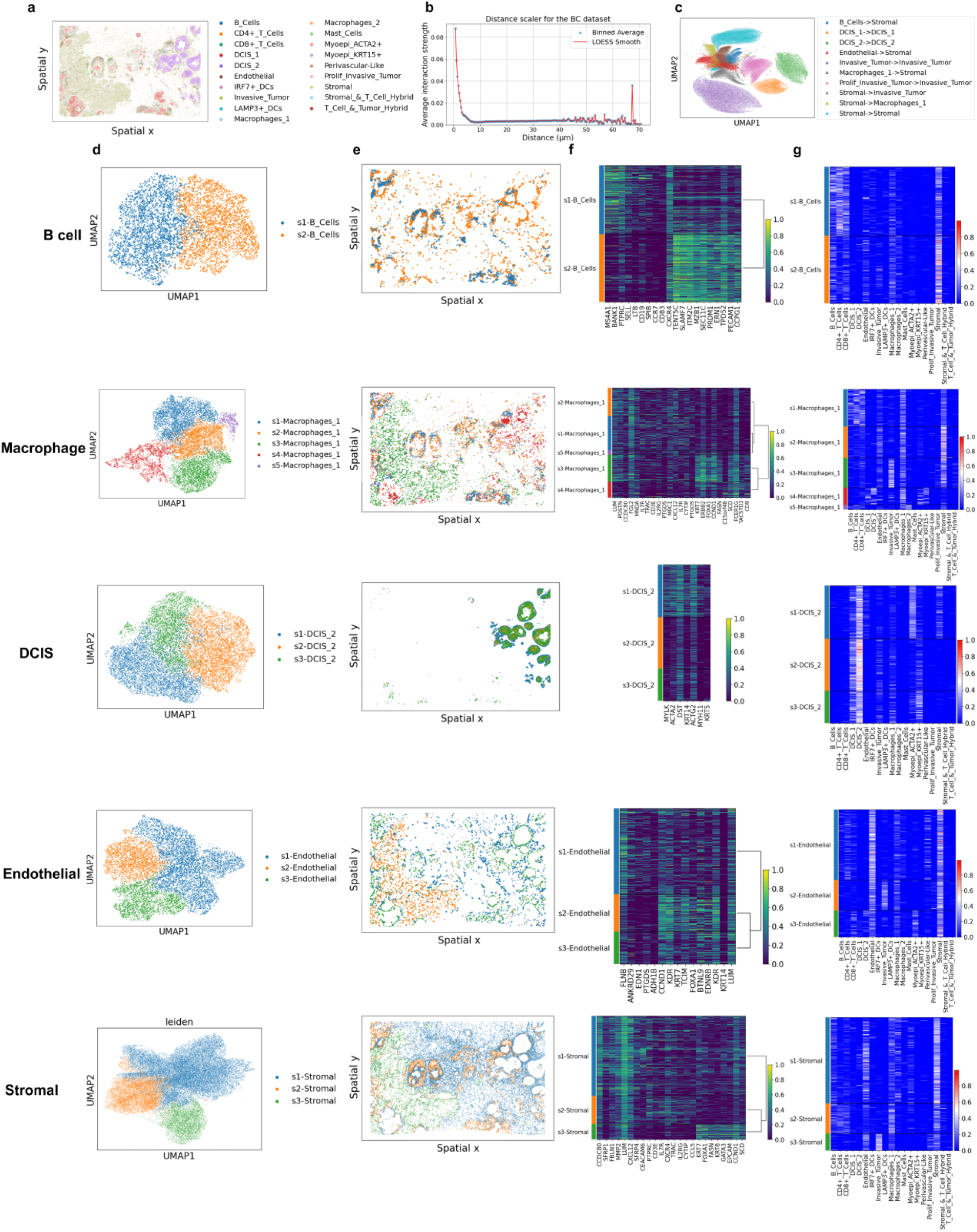
Analysis of CCI-driven within-cell-type heterogeneity in the breast cancer dataset. **a,** Spatial distribution of 19 cell types. **b,** The LOESS curve of the estimated CCI strength decreased with increasing distance between interacting cells. **c,** UMAP visualization of CCI pairs. **d,** UMAP visualization of B cell, macrophage, ductal carcinoma in situ (DCIS), endothelial cell, and stromal cell subgroups. **e,** Spatial distribution of B cell, macrophage, DCIS, endothelial cell, and stromal cell subgroups. **f,** Heatmap of the top differentially expressed genes in B cell, macrophage, DCIS, endothelial cell, and stromal cell subgroups. **g,** Neighboring cell type influence on B cell, macrophage, DCIS, endothelial cell, and stromal cell subgroups.

We then examined how immune, tumor, and structural cells in the TME are influenced by their neighborhood cells. CCI-informed microenvironmental features clustered B cells into two subgroups (Fig. 5d) with distinct transcriptional profiles (Fig. 5f). Subgroup 1 showed increased expression of *MS4A1*, *BANK1*, *PTPRC*, and *CD19*, and decreased expression of *TENT5C*, *SLAMF7*, *ITM2C*, and *NZB1* (adjusted 𝑝 < 1 × 10^−2^^4^), and was strongly influenced by CD4+ T cells (Fig. 5g). Macrophages were clustered into five subgroups (Fig. 5d). Subgroup 3, mainly influenced by stromal cells and invasive tumor cells (Fig. 5g), showed increased expression of *FOXA1*, suggesting that tumor-macrophage interactions may modulate immune responses in cancer^75^.

For tumor cells, we focused on a group of ductal carcinoma in situ (DCIS_2). These cells were clustered into three subgroups, where subgroup 1 located on the tumor boundary, and subgroups 2 and 3 were inside the tumor. Subgroup 1 showed increased expression of *MYLK*, *ACTA2*, and *KRT14* (adjusted 𝑝 < 1 × 10^−2^^4^), suggesting a progressive role for DCIS at the tumor boundary^76^. We also analyzed structural cells, including endothelial and stromal cells. Endothelial subgroup 2, strongly influenced by invasive tumor cells (Fig. 5g), showed upregulated expression of *CCND1*, *KDR*, *KRT7*, and *TCM* (adjusted 𝑝 < 1 × 10^−2^^4^), suggesting tumor-driven angiogenic signaling^77^. Stromal subgroup 3, also influenced by invasive tumors, showed increased expression of *KRT7*, *FOXA1*, *FASN*, and *KRT8* (adjusted 𝑝 < 1 × 10^−2^^4^), indicating that tumor-resident stromal cells contribute to tumor invasion by promoting the expression of basal epithelial markers^78^.

### Consistyency of CCI signals across datasets

To demonstrate the robustness of GITIII, we compared the identified interactions between cell types across datasets that differ by species and platform. In the mouse PMC dataset, GITIII identified interactions between L4/5 IT and L2/3 IT neurons that upregulate the *Rorb* expression in L2/3 IT neurons (Fig. 2i, 2j). Similar interactions were observed in the human SEA-AD dataset, where the interactions between L4 IT and L2/3 IT neurons were associated with the upregulation of *RORB* in L2/3 IT neurons (Fig. 3h, 3i). These interaction patterns were found in the L2/3 IT subgroup adjacent to layer 4 in both datasets (Fig. 2e, 3e). In addition, GITIII consistently identified interactions between tumor cells and a subgroup of macrophages, as well as between tumor cells and a subgroup of endothelial cells, in both the NSCLC and breast cancer datasets (Fig. 4f, 5g), further indicating its cross-dataset generalizability.

Additionally, we analyzed the composition of interacting cell types for each receiver cell type across the four ST datasets. Consistently across datasets, we observed that the gene expression of certain receiver cell types was influenced by multiple sender cell types, with contributions extending beyond interactions with neighboring cells of the same type (Supplementary Fig. 4).

### Benchmarking and sensitivity analysis

We benchmarked GITIII against nine existing CCI analysis tools, COMMOT^8^, HoloNet^10^, NCEM^30^, CellChat^22^, CellPhoneDB^21^, Giotto^9^, SOAPy^16^, SpaTalk^14^, and stLearn^12^, as well as two graph deep learning backbone models, using the four ST datasets. Details of the benchmarked methods are provided in Supplementary Note 7. Performance was evaluated at both single-cell and cell-type levels.

At the single-cell level, we compared GITIII with three methods that estimate CCI signals on receiver cells, COMMOT, HoloNet, and NCEM, as well as graph attention networks and graph transformer. Evaluation was based on the intracell-type variance explained by CCI. Fig. 6a shows that GITIII achieved the highest intracell-type variance explained by CCI across all four datasets. NCEM ranked second among existing CCI analysis methods. The improvements of GITIII over NCEM were 20%, 36%, 44%, and 201%, respectively.

**Fig. 6.**
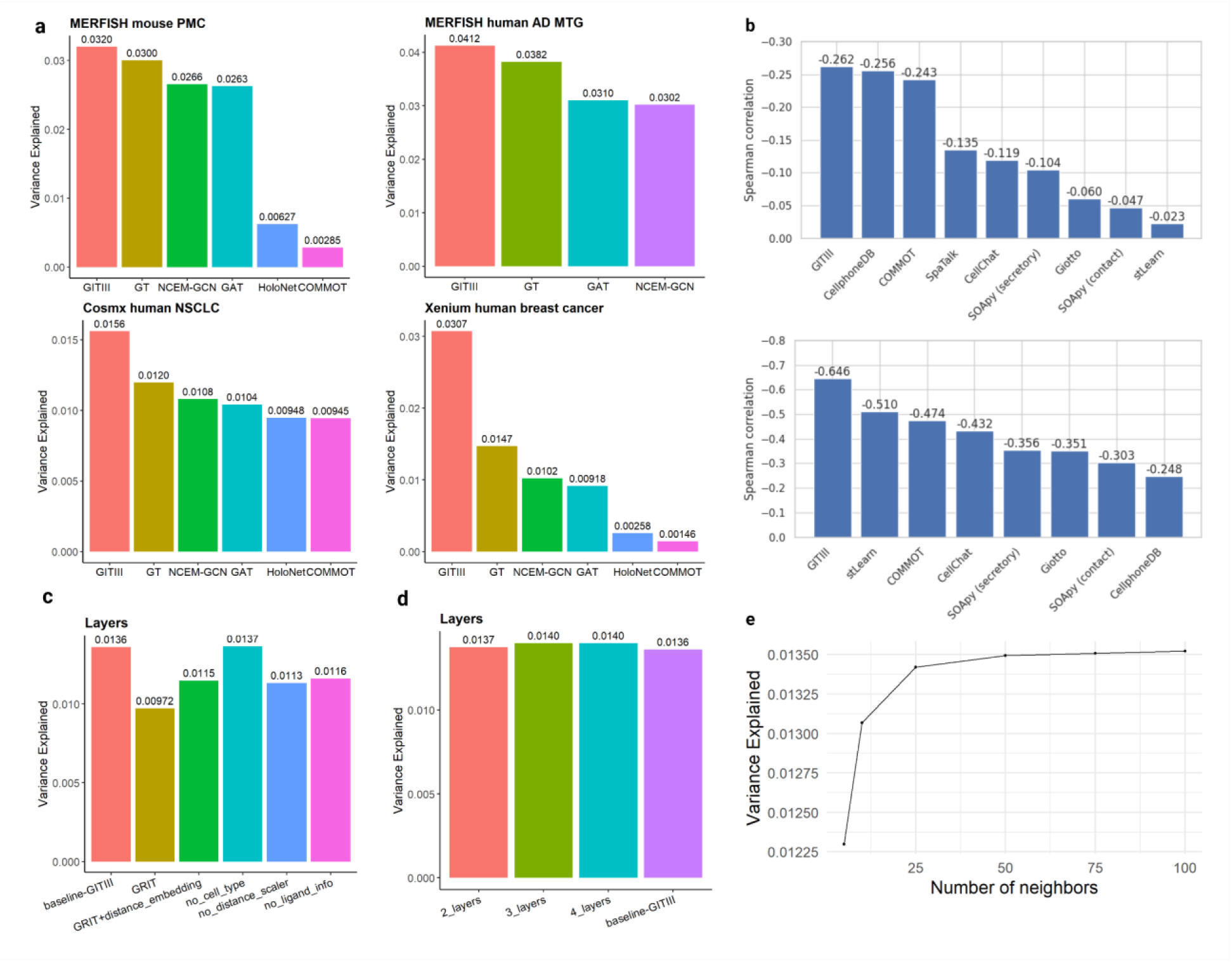
Benchmarking, ablation and sensitivity analysis. **a,** Single-cell level comparison of the intracell-type variance explained by CCI inferred from GITIII, COMMOT, HoloNet, NCEM, graph attention networks, and graph transformer in the four spatial transcriptomics datasets. **b,** Cell-type level comparison of the Spearman correlation coefficient between the aggregated CCI network strength and the Wasserstein distance between sender-receiver cell-type pairs inferred by GITIII, CellChat, CellPhoneDB, COMMOT, Giotto, SOAPy, SpaTalk, and stLearn in the NSCLC (upper) and breast cancer (lower) datasets. **c,** Comparison of the intracell-type variance explained by CCI in the ablation study, where the components or information removed or disabled include ligand expression in the sender cells, cell type information, distance scaler, and all modifications to the GRIT block, using the NSCLC dataset. **d,** Intracell-type variance explained by CCI inferred from GITIII using one to four encoder layers in the NSCLC dataset. **e,** Intracell-type variance explained by CCI inferred from GITIII using different numbers of cells in the cellular neighborhood in the NSCLC dataset.

At the cell-type level, we conducted benchmarking by comparing CCI networks inferred by GITIII, CellChat, CellPhoneDB, COMMOT, Giotto, SOAPy, SpaTalk, and stLearn. Performance was evaluated using the Spearman correlation coefficient (SCC) between the aggregated CCI network strength and the Wasserstein distance between sender-receiver cell-type pairs. A more negative SCC (i.e., larger absolute value) indicates better performance in capturing spatial CCI patterns. The two MERFISH datasets^33,45^, however, contain only 254 and 140 genes, respectively. Because all methods we compared relied on their built-in ligand-receptor databases, they identified fewer than 10 ligand-receptor pairs in these datasets, and in some cases none at all. Given the limited number of ligand-receptor pairs detected, we excluded these two datasets from the cell-type-level benchmarking. Fig. 6b shows that GITIII achieved the most negative SSC among all methods in both the NSCLC and breast cancer datasets.

To gain insights into the contribution of individual components to the performance of GITIII, we conducted an ablation study using the NSCLC CosMx dataset. The components or information removed or disabled include ligand expression in the sender cells, cell type information, distance scaler, and all modifications to the GRIT^79^ encoder on which our model is based. Among the individual ablations (Fig. 6c), the removal of all modifications to the GRIT block had the most impact on model performance, with the intracell-type variance explained by CCI decreased from 0.0136 to 0.00972, highlighting the importance of our modifications to the original GRIT model in encoding CCI. Removing each input information resulted in worse performance, except for cell type information. The intracell-type variance explained by CCI was 0.01361 when using cell type annotations and 0.01366 without them. This negligible difference suggests that the estimation of CCI influence on cell states in GITIII is robust to the absence of cell type annotations. GITIII’s architecture with attention calculation and feature embedding can learn cell type information from the input data to encode CCI. However, we acknowledge that the downstream interpretation analysis and visualization rely on cell type labels.

In the analysis of ST data, most GNN-based models employ multiple GNN layers, while GITIII uses a single-layer GRIT block to preserve interpretability and still outperforms other models. Fig. 6d shows that adding extra encoding layers only led to marginal improvement but would reduce model interpretability. Therefore, we used a single-layer graph transformer encoder to preserve interpretability. Finally, we evaluated the sensitivity of GITIII to the number of cells in the cellular neighborhood. The intracell-type variance explained by CCI increased as the number of neighboring cells increased and plateaued when the size of the cellular neighborhood reached 50 in the NSCLC dataset (Fig. 6e). To reduce computational resources, we set the size of the cellular neighborhood to 50.

To assess the robustness of GITIII to different quality of cell segmentation and marker quantification, we applied it to the breast cancer Xenium data processed by three cell segmentation methods, Voronoi, JSTA^80^, and BIDCell^81^. At the single-cell level, the spatial organization, gene expression, and interacting cell types within the local microenvironment of CCI-informed subgroups, for macrophages, fibroblasts, and SCGB2A2+ malignant cells, were broadly consistent across the three datasets (Supplementary Note 8, Supplementary Fig. 5-7). At the cell-type level, the composition of interacting cell types for each receiver cell type was also generally consistent across the three datasets (Supplementary Fig. 8). Together, these results demonstrate that GITIII is robust to differences in cell segmentation.

## Discussion

Imaging-based ST provides spatial gene expression patterns at single-cell resolution, enabling a deep understanding of intercellular communication critical for tissue development and organ function. Disruptions in normal CCI are implicated in disease onset and progression, such as neurodegeneration and cancer. We developed GITIII, a state-of-the-art GNN model that identifies single-cell-level interactions in imaging-based ST data. It integrates MLP-based distance encoding with physics-informed attention calculation, outperforming existing methods in capturing the complex spatial patterns of CCI. GITIII’s interpretable architecture allows estimation of the influence of sender cells on the gene expression of receiver cells, facilitating downstream analyses including visualization of spatial CCI patterns, CCI-informed cell clustering, identification of significant cell-type interactions, CCI network analysis, and comparison of CCI strength across conditions. We applied GITIII to four ST datasets across multiple species, organs, and platforms, demonstrating superior performance over existing GNN models in decoding CCI influences. These applications advanced our understanding of brain structure and the TME.

Despite its advantages, GITIII has several limitations. First, it is designed for imaging-based ST platforms that measure only a limited set of genes, often with sparse or missing ligand-receptor pair information. As a result, the downstream signaling pathways representing functional consequences of CCI remain unexplored. To address this, we plan to integrate scRNA-seq and ST data in future work to characterize the functional influence of CCI. Second, the inferred CCI only indicate the association between cell states and their niche, not a causal interpretation. Thus, the biological results generated by our method require further validation through experimental studies. Third, signal spillover between segmented neighboring cells is a known issue in imaging-based ST, often arising from the imperfect cell segmentation. Such artefacts can confound and complicate the identification of cell types and degrade data quality, thereby affecting the inference of cellular interactions.

## Methods

### Data preprocessing

Consider an imaging-based single-cell ST dataset that includes a cell-by-gene expression matrix, spatial coordinates of cells, and cell type annotations. We first log-transformed the normalized expression matrix. Then, we computed a distance matrix using the Euclidean distance between cells. To ensure data quality, we excluded cells that had no neighboring cells within a radius of 𝑟 pixels (default is 80). These cells, along with their corresponding rows in the expression matrix and both rows and columns in the distance matrix, were removed. Lastly, for each cell type, we calculated the cell-type expression by averaging the gene expression levels across all cells of that type. The cell-state expression of each cell was then calculated as the deviation of its gene expression from the average expression of its respectively cell type.

### GITIII algorithm

The GITIII pipeline consists of four components: a deep learning model with an embedding module and a modified graph inductive bias transformer to estimate the influences of sender cells on the gene expression of receiver cells; a CCI pattern visualization module; a cell clustering analysis module; and a network analysis module. Fig. 1 shows the schematic diagram of GITIII. For convenience, a list of notations used below is included in Supplementary Note 1.

### Deep-learning model architecture

GITIII uses a lightweight, self-supervised, single-layer graph transformer-based language model to predict the cell-state expression of each cell based on the spatial location and gene expression profiles of its neighboring cells, thereby learning how neighboring sender cells of varying types and states interact with a central receiver cell (Fig. 1a). The ST data can be represented as a spatial graph, where nodes are cells and edges are spatial proximity between cells. For each cell, GITIII first constructs a cellular neighborhood by identifying its (𝑘 − 1) nearest neighboring cells to form a subgraph. This process converts the original spatial graph into 𝑁 graphs of size 𝑘, where 𝑁 is the total number of cells in the ST data, and each graph represents a cellular neighborhood. Conceptually, these cellular neighborhoods are treated as sentences, with each cell representing a word. GITIII processes these subgraphs using a deep learning model consisting of two parts: an embedding module (Fig. 1b) and a single-layer modified graph inductive bias transformer encoder (Fig. 1c). The model outputs an influence tensor that quantifies the impact of each neighboring sender cell on the gene expression of the central receiver cell.

Both CCI within the cellular neighborhood and intracellular biological processes, such as cell differentiation, influence cell states. GITIII is not designed to fully explain cell-state variation, but to capture the component attributable to CCI influence from neighboring cells. The model is trained by minimizing the sum of squared differences between the observed and predicted cell-state expression of receiver cells, where the predicted expression is defined as the aggregated CCI influence from neighboring cells. Importantly, the aggregated CCI influence estimated by GITIII is not intended as a full prediction of cell-state expression, but rather as a way to quantify the influence from neighboring cells.

### Model input

The input of GITIII includes three types of information from a cellular neighborhood: (1) the cell-by-gene expression matrix, denoted as Exp ∈ ℝ^𝑘×𝑐^, where 𝑐 is the number of genes measured in the ST data; (2) the distance matrix, denoted as 𝐷 ∈ ℝ^𝑘×𝑘^, the Euclidean distance between cells; and (3) the cell type vector, denoted as 𝑇 ∈ ℝ^𝑘^, representing cell type annotations of cells.

### Node feature embedding

The node feature embedding is generated in three steps: expression masking, expression embedding, and cell type embedding. Since GITIII infers CCI by predicting the cell-state expression of a central cell based on the information of its neighboring cells, and cells of the same type in close proximity often share CCI patterns, thus having similar cell-state expression (the closer the more similar), we mask the cell-state expression of neighboring cells of the same type as the central cell to prevent perfect prediction. Specifically, within a cellular neighborhood of 𝑘 cells, where the first row in the cell-by-gene expression matrix, Exp ∈ ℝ^𝑘×𝑐^, corresponds to the gene expression profile of the central cell 𝑐_1_. For each neighboring cell 𝑐_𝑖_, if it is of the same cell type as the central cell, its gene expression is replaced by its cell-type expression; otherwise, its observed gene expression is retained. Formally, it is defined as

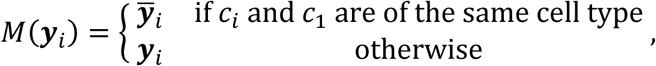

where 𝒚_𝑖_is the gene expression of 𝑐_𝑖_, and 𝒚̅_𝑖_denotes the cell-type expression of 𝑐_𝑖_, the average gene expression of all cells of the same type as 𝑐_𝑖_. Then, we apply a multilayer perceptron (MLP) to encode the expression features, producing the expression embedding. Finally, we incorporate cell type information by adding cell type embeddings to the node features. The node embedding

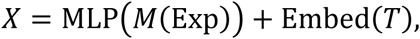

where 𝐷_𝑉_ is the dimension of the node embedding and Embed uses an embedding layer that maps discrete cell type indices to continuous vectors of fixed dimension, analogous to token embeddings commonly used in natural language processing.

### Distance embeddings

Given that the interaction between two cells is influenced by their distance, we model the effect of distance on CCI strength using five distance-based features, which differ from the commonly used exponential decay diffusion model, to better describe dynamic processes in biological systems. We then apply MLPs to these features. Specifically, we generate two distance embeddings to be used in edge embeddings and one to be used in the graph transformer encoder. Of these three distance embeddings, two are scaled to a range of 0 to 1 using the sigmoid function.

The design and rationale behind these distance embeddings are explained in detail in Supplementary Note 2.

### Edge feature embedding

Edge features are constructed to capture the distance between interacting cells and the effects of ligands from the sender cell on the receiver cell. The ligand effect is modeled as the ligand expression in the sender cell, scaled by one of the distance embeddings. Briefly, we first multiply the ligand expression by the scaled distance embedding and apply a linear transformation. Then, we incorporate the spatial organization by concatenating the resulting tensor with an unscaled distance embedding and apply another linear transformation to produce the final edge embedding, denoted as 𝐸 ∈ ℝ^𝑘×𝑘×𝐷𝐸^, where 𝐷_𝐸_ is the dimension of the edge embedding. Further details on the edge embedding are provided in Supplementary Note 3.

### Graph attention calculation

We develop a graph attention method based on the Graph Inductive bias Transformer (GRIT)^79^ to capture the complexity of CCI. GRIT is a state-of-the-art graph transformer that does not rely on message-passing. Instead, it incorporates inductive biases through learned relative positional encodings based on random walk probabilities. GRIT encodes node and edge features by integrating position information, node features, and edge features using attention mechanisms. This architecture is particularly well suited for modeling CCI, where gene expression of cells (node features), spatial organization (positional information), and ligand expression in sender cells (edge features) interact with each other and collectively influence the gene expression of receiver cells. To adapt GRIT for encoding cellular neighborhood information in ST, we implemented three main modifications in GITIII: (a) we replaced the random walk graph structure encoding designed for sparse graphs with distance embeddings to better model the dense, two-dimensional spatial organization of cellular neighborhoods; (b) we incorporated a distance scaler into the attention mechanism to explicitly model the decay of CCI effects with increasing intercellular distance; and (c) we removed the degree scaler, which adjusts for variation in node connectivity, as the cellular neighborhoods in our setting have uniform connectivity by construction. These changes tailor the GRIT framework to the unique spatial and biological characteristics of single-cell ST data and improve its capacity to capture biologically meaningful CCI. In addition, we refine the attention calculation by computing scores only from neighboring sender cells to a central receiver cell and performing a single pass of the attention calculation. This ensures that the aggregated values are solely determined by the features of individual cells, avoiding multi-step contributions from indirect sources. Following is a summary of the key ideas in our graph attention calculation, with a detailed description of the algorithm provided in Supplementary Note 4.

We first construct the Key, Query, and Value matrices using three linear transformations of the node features. The attention matrix is then computed by multiplying the Key and Query matrices to capture how signaling pathway genes, cell type, and other regulatory genes influence the CCI effects. Next, we update the edge features by integrating the edge embeddings with the unscaled distance embeddings and the attention matrix, thereby encoding distance, gene expression, and cell type characteristics relevant to CCI. A linear transformation is then applied to the updated edge features to generate the initial raw attention scores. We discard the self-attention score of the receiver cell and apply a softmax function for normalization. Finally, we update the Value matrix by integrating it with the edge features whose destination is the receiver cell using an MLP. This operation models interactions between edge and node features to enhance the graph representation and encoding capacity.

The Value matrix 𝑉 = 𝑉[1:, :] ∈ ℝ^(𝑘−1)×𝐷𝐸^ (subscripts starting from 0) represents the attention values of all neighboring cells, where the attention scores from the neighboring sender cells to the receiver cell are denoted as 𝛼 ∈ ℝ^1×(𝑘−1)^ . We compute the influence matrix 𝑖 ∈ ℝ^(𝑘−1)×𝑐^ by multiplying the Value matrix with the attention scores and applying an MLP to transform the last dimension from 𝐷_𝑉_to the number of measured genes 𝑐. Since each row of the influence matrix is only determined by a cell’s type, gene expression features, and its distance to the receiver cell, we interpret it as representing the influence of the (𝑘 − 1) neighboring cells on the expression levels of 𝑐 genes in the central cell. Summing over the first dimension of 𝑖 yields the predicted cell-state expression 𝑦^ ∈ ℝ^𝑐^, which is then compared to the observed cell-state expression to infer how neighboring sender cells influence the central receiver cell.

### Interpretation of the influence tensor

We construct the influence tensor, denoted as 𝐼 ∈ ℝ^𝑁×(𝑘−1)×𝑐^, by concatenating the influence matrices learned from each subgraph representing a cellular neighborhood. Since all non-attention-based transformations in GITIII are performed along the last dimension of the input tensor, and the attention mechanism preserves both the order and representation of each token (cell), the identity of each cell remains intact throughout the encoding process. Thus, the entries in the influence tensor can be interpreted as the impact of the (𝑘 − 1) neighboring cells on the expression levels of 𝑐 genes in the central cell. Instead of using attention scores alone to quantify the influence of CCI on receiver-cell expression, the entries in the influence tensor, calculated as the product of the embedding vector and the corresponding attention score, represent the contributions of neighboring cells to the cell-state expression of the central receiver cell. The justification for using the influence tensor as a measure of CCI influence is presented in Supplementary Note 5.

### Normalization and aggregation of the influence tensor

To compare the influence of neighboring cells on the gene expression of a central cell within a cellular neighborhood, we calculate a proportional influence tensor, denoted as 𝐼_𝑝_ ∈ ℝ^𝑁×(𝑘−1)×𝑐^, by normalizing the influence tensor 𝐼. Specifically, for each gene in a given cell, we divide the influence values from its neighboring cells by the sum of the absolute values of the influences on that gene across all neighboring cells. If the absolute values of the proportional influence are below a threshold (default is 0.02), we discard these subtle influences by setting the corresponding entries of 𝐼 to 0, and then recalculate 𝐼_𝑝_ as described above. The default threshold of 0.02 was chosen to ensure that removing weak interactions has minimal impact on the intracell-type variance explained by CCI. Across all four datasets analyzed, applying this threshold led to a reduction of less than 5% in the intracell-type variance explained, supporting its use as a conservative and robust cutoff. Additionally, we compute the average of the absolute proportional influence values across all genes to estimate the overall strength of CCI between two interacting cells. The resulting matrix, denoted as 𝐼_𝑠_ ∈ ℝ^𝑁×(𝑘−1)^, serves as a metric for identifying significant CCI pairs in downstream analysis.

To estimate the effect of different sender cell types on the gene expression of receiver cells, we aggregate the influence values by sender cell type within each cellular neighborhood, resulting in an aggregated influence tensor, 𝕀 ∈ ℝ^𝑁×𝑡×𝑐^, where 𝑡 is the number of cell types in the ST data, and the entries represent the impact of neighboring cell types on the expression levels of 𝑐 genes in the central cell. If specific cell types are absent among the neighboring cells in a cellular neighborhood, the corresponding entries are set to 0. Similarly, we calculate a proportional aggregated influence tensor, 𝕀_𝑝_ ∈ ℝ^𝑁×𝑡×𝑐^, by normalizing the aggregated influence tensor. For each gene in a given cell, we normalize the aggregated influence values by the sum of the absolute values of the aggregated influences on that gene across all neighboring cell types.

### Visualization of CCI patterns

The CCI pattern visualization module in GITIII offers three types of visualizations to understand the spatial patterns and heterogeneity of CCI, including (1) visualization of the relationship between CCI strength and the distance between interacting cells, (2) visualization of CCI pairs, and (3) visualization of the spatial distribution of CCI.

To visualize the relationship between CCI strength and the distance between interacting cells, we flatten the influence tensor 𝐼 and plot the absolute value of each entry against the distance between the corresponding sender and receiver cells. A locally estimated scatterplot smoothing (LOESS) curve is fitted to demonstrate the decay of CCI strength with increasing distance between interacting cells. For the visualization of CCI pairs, we treat each cell on the slide as a receiver cell and select the top five sender cells from its neighborhood with the strongest interactions based on the overall strength values in 𝐼_𝑠_to form CCI pairs. These selected CCI pairs are visualized on a UMAP using the estimated influence values on the cell-state expression of the corresponding receiver cell as features, where each triangle represents a CCI pair. Finally, to visualize the spatial distribution of CCI influencing the expression level of a specific gene, we focus on the top 5% of CCI pairs with the largest absolute influence values for that gene among the selected CCI pairs across all cellular neighborhoods. The arrow connecting the corresponding sender and receiver cells is drawn on a spatial map, with the color of each arrow representing the influence value of the interaction.

### Cell clustering analysis

The cell clustering analysis module in GITIII provides a quantitative interpretation of the local microenvironment and its impact on cell-state expression, including (1) clustering cells into subgroups based on CCI-informed microenvironmental features, (2) spatial visualization of cell subgroups, (3) differential expression analysis across subgroups, and (4) investigation of microenvironmental influence on cell-state expression.

To perform cell clustering within each cell type, we first construct CCI-informed microenvironmental features by flattening the last two dimensions of the aggregated influence tensor 𝕀 into a feature matrix 𝐹 ∈ ℝ^𝑁×(𝑡𝑐)^. We then apply the Leiden algorithm^82^ to cluster cells into subgroups based on these microenvironmental features and visualize cell subgroups using UMAP. The distinction among these subgroups reflects differences in their surrounding cellular neighborhood, rather than differences in gene expression alone. We expect that CCI-informed subgroups mainly differ in their local microenvironment and tend to locate in spatially distinct regions.

For each cell type, we analyze the differences among its subgroups by examining their spatial organization, gene expression, and interacting cell types within the local microenvironment. Specifically, we visualize the spatial locations of subgroups on a spatial map. Next, we perform differential expression analysis to identify the top marker genes for each subgroup using the Wilcoxon rank sum test. To assess the influence of neighboring cell types on the expression level of a specific gene in receiver cells of different subgroups, we visualize the aggregated influence values of the gene of interest in a heatmap. Additionally, we compute the average of the absolute proportional aggregated influence values across all genes and visualize them in a heatmap to better understand the overall influence of neighboring cell types on receiver cells of different subgroups. These analyses generate a comprehensive view of the contributions of interacting cell types within the local microenvironment that influence cell-state expression across subgroups within each cell type.

### Construction of CCI network

If there is a significant interaction between two cell types that influences the gene expression of a receiver cell type, we expect that the proportional influence value is correlated with the cell-state expression of that gene among CCI pairs of these two cell types. To identify such significant interactions, for a fixed gene 𝑙 and a pair of cell types, say sender cell type 𝐴 and receiver cell type 𝐵, we assess the correlation between the proportional aggregated influence values of neighboring cell type 𝐴 on gene 𝑙 in receiver cells of type 𝐵 and the cell-state expression of gene 𝑙 in these receiver cells, using the Pearson correlation coefficient. Pearson correlation analysis is performed across all genes and all pairs of cell types, with the corresponding z-scores calculated. To account for multiple hypothesis testing, significance is adjusted using the Benjamini-Hochberg false discovery rate (FDR) control. Sender cell type 𝐴 is considered to significantly influence the expression level of gene 𝑙 in receiver cell type 𝐵 if the FDR-adjusted p-value < 0.05. For interactions that do not reach significance, we set their z-scores to 0. For each gene, these significant interactions between cell-type pairs are used to construct a CCI network that illustrates how cell types influence the expression of that gene.

When multiple slides are available, we aggregate the individual CCI networks across slides using Stouffer’s method to compute the combined z-scores and corresponding p-values.

To evaluate the composition of interacting cell types for each receiver cell type, we first calculate the interaction strength between each sender-receiver cell-type pair, defined as the average of the absolute proportional aggregated influence values across all genes and all receiver cells of the same type. We then normalize these interaction strength by the sum of the interaction strength across all sender cell types, illustrating the relative contributions of different interacting cell types for each receiver cell type.

### Comparison of CCI network strength between conditions

To quantify CCI network strength, we introduce a microenvironment-adjusted network strength metric to facilitate comparisons across slides. This metric is less susceptible to batch effects compared to the traditional network strength, which is commonly defined as the product of the mean ligand and receptor expression levels^1^. For a fixed gene 𝑙 and a pair of cell types, say sender cell type 𝐴 and receiver cell type 𝐵, we consider the proportional aggregated influence value for each receiver cell of type 𝐵 from the corresponding entries in 𝕀_𝑝_. To adjust for the local microenvironment, we subtract 𝑛_𝐴_/(𝑘 − 1), where 𝑛_𝐴_ is the number of neighboring cells of type 𝐴. This adjustment accounts for the cell-type composition of the cellular neighborhood. We then aggregate these microenvironment-adjusted proportional aggregated influence values by receiver cell type within each slide to obtain the CCI network strength for that sample. To assess the group-level CCI network strength, we compute the average CCI network strength across all samples within the group. To compare CCI network strength between conditions, we use the Wilcoxon rank sum test, with significance levels adjusted using the FDR to account for multiple hypothesis testing. Interacting cell-type pairs are considered to have significantly different CCI network strengths between conditions if FDR < 0.1.

### Summary of the advantages of GITIII

The main advantages of GITIII over existing GNN-based CCI prediction methods are summarized as:

1. GITIII is designed to estimate the effects of neighboring sender cells on the gene expression of receiver cells through cell-cell interactions, which is a key aspect of CCI inference that has not been explicitly studied by other GNN-based methods.
2. While existing GNN-based methods rely on the availability of ligand-receptor pairs measured in ST data, GITIII can be applied to imaging-based ST datasets, where only a limited number of genes are measured and the covered ligand-receptor pairs are sparse or absent.
3. Most existing GNN-based CCI analysis methods consider 2D ST data as sparse graphs, potentially losing interaction information due to limited pairwise distances between cells. In contrast, GITIII models ST data as dense graphs and uses an expressive graph transformer to better encode spatial CCI patterns. Our benchmarking results show that GITIII consistently outperformed other GNN-based methods in explaining the intracell-type variance.
4. GNN-based CCI prediction models use multiple layers, with each layer repeatedly propagating information from neighboring nodes to a central node. This multi-layer propagation makes it challenging to explicitly quantify the influence of individual neighboring cells on a central receiver cell. GITIII employs a single-layer graph transformer to directly calculate the influence from neighboring sender cells to the central receiver cell. To preserve the interpretability of the influence, GITIII does not apply any nonlinear transformation after aggregation of neighborhood information.
5. GITIII incorporates statistical analysis to construct CCI networks at the cell-type level and identify sender cell types that significantly influence the gene expression in receiver cell types. This CCI network inference has not been studied by other GNN-based CCI prediction methods.

### Implementation details

Data preprocessing was performed using Scanpy^83^, and the deep learning model was implemented in Pytorch 2.0.1. To balance model performance and computational efficiency, we selected a modest set of hyperparameters without the need for extensive GPU resources. Across all four ST datasets, the number of cells in a cellular neighborhood was set to 50 and the embedding dimension of the node features was fixed at 256. The dimensionality of the edge features was dertermined by the number of ligand genes measured in each dataset, resulting in the edge dimensions of 48 for the mouse PMC and SEA-AD datasets and 128 for the NSCLC and breast cancer datasets. We used the AdamW optimizer with a learning rate of 0.0001. A batch size of 256 was used, which required approximately 8 GB of GPU memory, for efficient model training on moderately powered hardware.

For all experiments, each dataset was randomly split into 80% training and 20% validation sets using a fixed random seed. Models were trained for 50 epochs, and the one with the smallest mean squared error on the validation set was selected for downstream analyses.

It took approximately one hour for GITIII to analyze a dataset containing 366,272 cells and 140 genes on a system with an NVIDIA A5000 GPU (24 GB VRAM) and four Intel Xeon CPUs with 100 GB of RAM. The runtime scales linearly with both the number of cells and the number of genes in the dataset.

### Variance decomposition into inter- and intra-cell-type variance

For each gene 𝑗, the total variance of its expression can be decomposed into intercell-type variance and intracell-type variance:

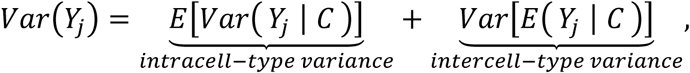

where 𝐶 is cell type. The first term represents the variation of gene expression within cell types (intracell-type variance), corresponding to differences in cell state, and the second term represents the variation of gene expression across cell types (intercell-type variance). A gene with a higher proportion of intracell-type variance relative to the total variance 𝑉𝑎𝑟(𝑌_𝑗_) is more strongly associated with cell state, whereas one with a higher proportion of intercell-type variance is more strongly associated with cell type. It is important to note that a high proportion of cell-state variation does not necessarily imply that a gene’s expression is influenced by CCI. Cell-state expression can also be influenced by intracellular biological processes, which are not captured by our model.

### Benchmarking metrics Single-cell-level evaluation

Since CCI influence cell states and contribute to intracell-type variability, we use the intracell-type variance explained by CCI to evaluate method performance at the single-cell resolution. Specifically, we use the following metric:

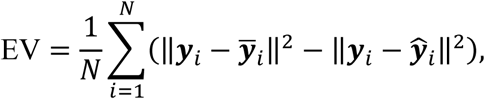

where 𝒚_𝑖_is the observed gene expression of cell 𝑖, 𝒚̅_𝑖_is the cell-type expression of cell 𝑖, the average gene expression of all cells of the same type as cell 𝑖, and 𝒚^_𝑖_ is the model-predicted expression. This metric, also used in NCEM^30^, captures the improvement in predictive accuracy by modeling CCI over simply using the cell-type average expression, and quantifies how much variability in cell states is explained by CCI effects.

### Cell-type-level evaluation

We evaluate method performance by comparing the inferred CCI nextworks at the cell-type level. The evaluation metrics based on significant ligand-receptor pairs are not applicable to GITIII, because it neither uses the receptor information nor identifies significant ligand-receptor interactions. Instead, we evaluate performance by computing the Spearman correlation coefficient (SCC) between the aggregated CCI network strength and the Wasserstein distance between sender-receiver cell-type pairs. It is commonly assumed that spatially adjacent cell types are more likely to interact than distant ones^4,18^. A more negative SCC (i.e., larger absolute value) indicates better performance in capturing spatial CCI patterns.

### Evaluation of cell segmentation on GITIII’s performance

To assess the robustness of GITIII to differences in cell segmentation, we considered three methods: (1) classical Voronoi expansion, (2) JSTA^80^, a joint cell segmentation and cell type annotation approach based on an extension of the watershed algorithm, and (3) BIDCell^81^, a biologically-informed deep learning approach. After applying cell segmentations to the breast cancer Xenium dataset, cell type annotations were assigned using scClassify^84^, with detailed processing steps described previously^81^. Because the number and labels of cell types were consistent across the three segmentation datasets, but differed from those in the original dataset^26^, we focused our evaluation on the consistency of CCI results inferred by GITIII across the three datasets. In the first replicate slide from sample 1, Voronoi, JSTA and BIDCell produced 106,227, 107,131, and 103,209 cells, respectively.

### Datasets

The mouse primary motor cortex (PMC) ST dataset generated by MERFISH^33^ contains 254 genes measured in 284,098 cells across 64 slices from two mice, with annotations of 24 cell types. We analyzed slice 201 from mouse 1 with 6,137 cells. Among the 254 genes, 47 were identified as ligand genes and no ligand-receptor pairs were found in the ligand-receptor databases CellChatDB^22^ and NeuronChatDB^6^.

The SEA-AD ST dataset generated by MERFISH^45^ contains 140 genes measured in 366,272 cells across 69 slides of the human middle temporal gyrus (MTG) from 27 donors. Sample ‘H20.33.001.CX28.MTG.02.007.1.02.03’ with 15,225 cells from 24 cell types was used for demonstration. We also analyzed all 69 slides and aggregated CCI networks for group comparison.

Among the 140 genes, 28 were identified as ligand genes and 2 ligand-receptor pairs were found in CellChatDB^22^ and NeuronChatDB^6^.

The human non-small cell lung cancer (NSCLC) ST dataset generated by CosMx^25^ includes 980 genes measured in 769,114 cells across eight formalin-fixed paraffin-embedded (FFPE) slides from five NSCLC tumors, with annotations of 18 cell types. We analyzed sample ‘Lung 6’ with 89,949 cells.

The human breast cancer ST dataset generated by Xenium^26^ contains 313 genes measured across three slides. Two replicates of 5 μm sections from sample 1, which include 274,492 cells from 19 annotated cell types, were analyzed. Among the 313 genes, 33 were identified as ligand genes.

### Data availability

The mouse PMC MERFISH dataset is available at ftp://download.brainimagelibrary.org:8811/02/26/02265ddb0dae51de/. The SEA-AD MERFISH dataset is available at https://sea-ad-spatial-transcriptomics.s3.amazonaws.com/middle-temporal-gyrus/all_donors-h5ad/SEAAD_MTG_MERFISH.2024-02-13.h5ad. The NSCLC CosMx dataset is available at https://nanostring.com/products/cosmx-spatial-molecular-imager/ffpe-dataset/nsclc-ffpe-dataset/. The breast cancer Xenium dataset is available at https://www.10xgenomics.com/products/xenium-in-situ/preview-dataset-human-breast. The ligand-receptor databases of CellChat and NeuronChat are publicly available, and an integrated version is available on our GitHub page.

## Code availability

GITIII is implemented in open-source Python using PyTorch. The source code and a detailed tutorial are freely available at https://github.com/lugia-xiao/GITIII. The Code to reproduce the results in the manuscript are available at https://github.com/lugia-xiao/GITIII_reproducible.

## Supporting information

https://drive.google.com/file/d/1SJgG6oOVC1OgxTLeYypxDwqpb7-3LzAJ/view?usp=sharing

## Acknowledgments

This study was supported by the National Institutes of Health (NIH) grant R01LM014087. We are grateful to Drs. Jean Y.H. Yang and Yingxin Lin for generously sharing their data generated using different cell segmentation methods.

## Author contributions

X.X. and Z.W. conceptualized the study. X.X. designed and implemented the model algorithm, analyzed the spatial transcriptomics data, and created the figures. L.Z., H.Z., and Z.W. helped refine the idea and analyses and aided result interpretation. X.X. and Z.W. wrote the manuscript Z.W. supervised the project. All authors read and approved the final manuscript.

---

**Extended Data Fig. 1.**
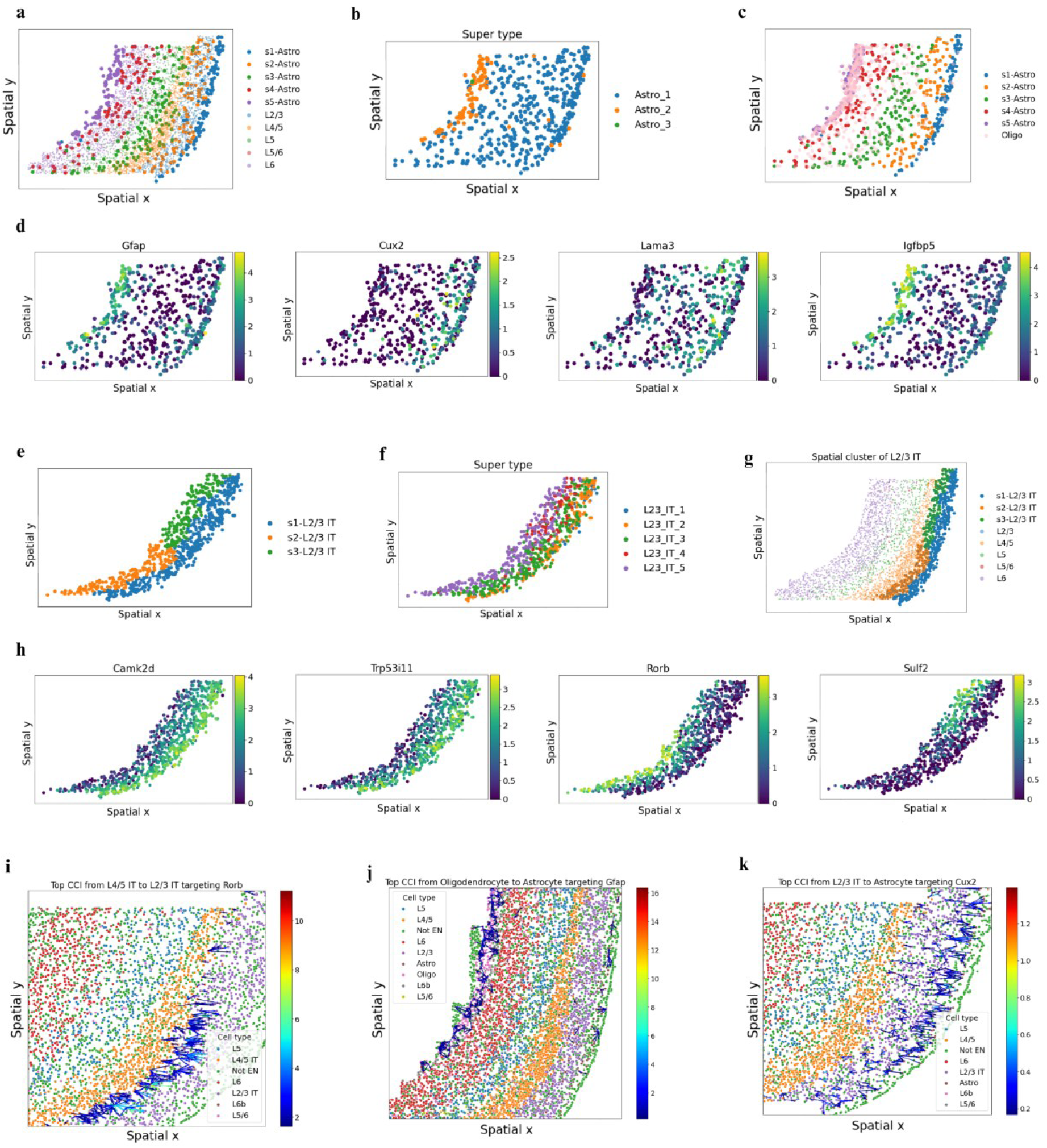
Spatial gene expression in subgroups identified by GITIII and spatial distribution of CCI influencing *Rorb*, *Gfap*, and *Cux2* in the mouse PMC datasets. **a,** Spatial distribution of astrocyte subgroups and excitatory neurons. Specifically, astrocyte subgroup 1 is located in layer 1, subgroup 2 is located in layers 2 and 3, subgroup 3 is located in layers 4 and 5, subgroup 4 is located in layer 6, and subgroup 5 is located in the white matter. **b,** Super type annotation of astrocytes by Zhang *et al.* **c,** Spatial distribution of astrocyte subgroups and oligodendrocytes. **d,** Spatial expression patterns of *Gfap*, *Cux2*, *Lama3*, and *Igfbp5* in astrocytes. Astrocyte subgroup 5 located in the white matter are physically close to oligodendrocytes and had higher expression of *Gfap* and *Igfbp5* compared to other astrocytes. In contrast, astrocyte subgroup 2 located in layers 2 and 3 are physically close to L2/3 IT neurons and had higher expression of *Cux2* and *Lama3*. **e,** Spatial distribution of L2/3 IT subgroups. Specifically, subgroup 1 is located in the upper region of layers 2 and 3, closer to layer 1, while subgroups 2 and 3 are located in the deeper region of layers 2 and 3, closer to layer 4. **f,** Super type annotation of L2/3 IT neurons by Zhang *et al.* **g,** Spatial distribution of L2/3 IT subgroups and other excitatory neurons. **h,** Spatial expression patterns of Camk2d, *Trp53i11*, *Rorb*, and *Sulf2* in L2/3 IT neurons. L2/3 IT subgroups 2 and 3 located in the deeper region of layers 2 and 3 are physically close to L4/5 IT neurons and had higher expression of *Rorb* and *Sulf2*, respectively, compared to other L2/3 IT cells. **i,** Spatial distribution of interactions from L4/5 IT to L2/3 IT neurons influencing the expression of *Rorb*. **j,** Spatial distribution of interactions from oligodendrocytes to astrocytes influencing the expression of *Gfap*. **k,** Spatial distribution of interactions from L2/3 IT neurons to astrocytes influencing the expression of *Cux2*.

**Extended Data Fig. 2.**
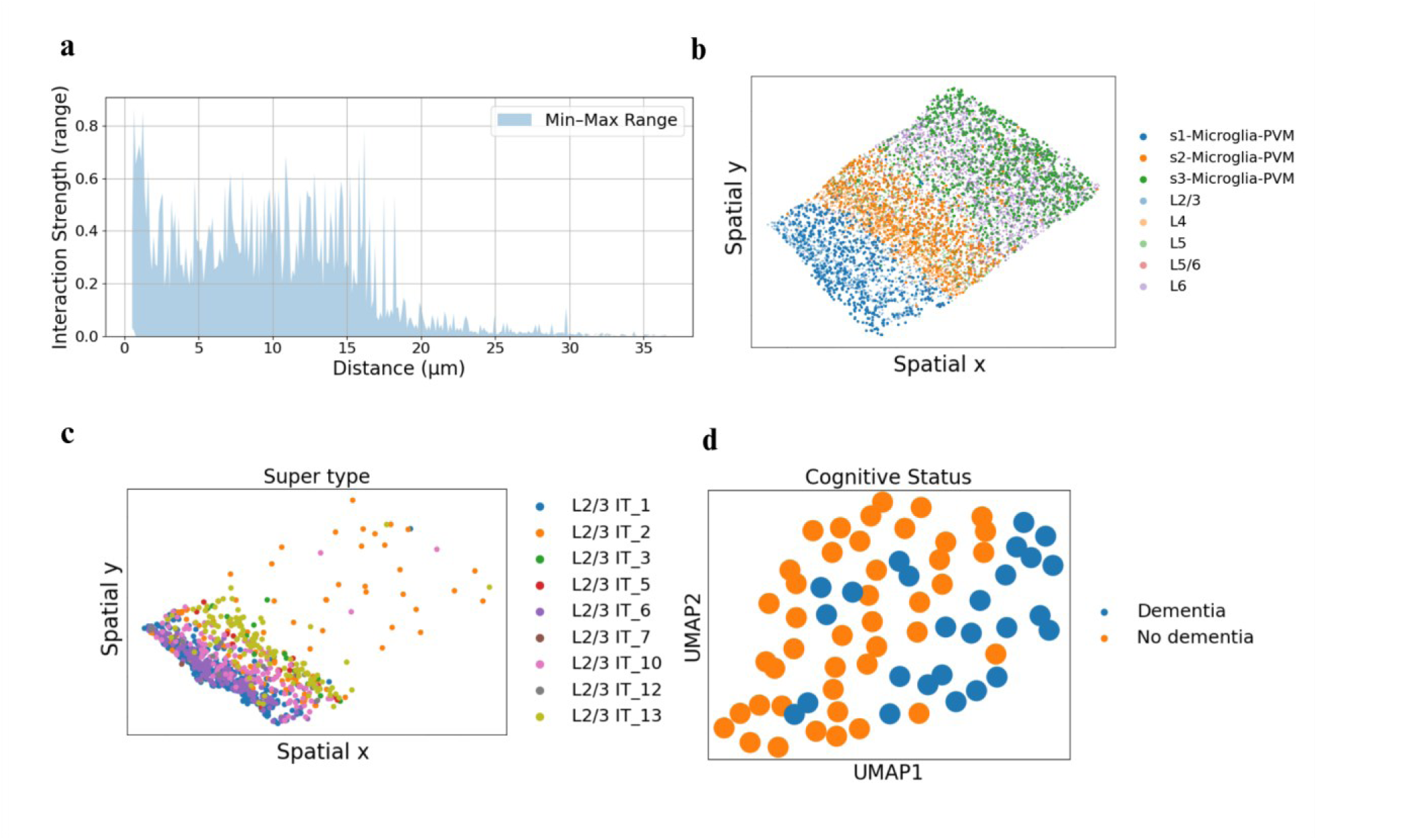
Spatial distribution of Microglia and L2/3 IT neurons in the SEA-AD dataset. **a,** The strength of interactions influencing the expression of *RORB*. **b,** Saptial distribution of microglia subgroups and excitatory neurons. Specifically, microglia subgroup 1 is located in layers 2 and 3, subgroup 2 in layers 4 and 5, and subgroup 3 in layer 6. **c,** Super type annotation of L2/3 IT neurons in Gabitto *et al.* **d,** UMAP visualization of slides using the inferred network strength as features, with color indicating slide from patients with or without dementia.

**Supplementary Fig. 1.**
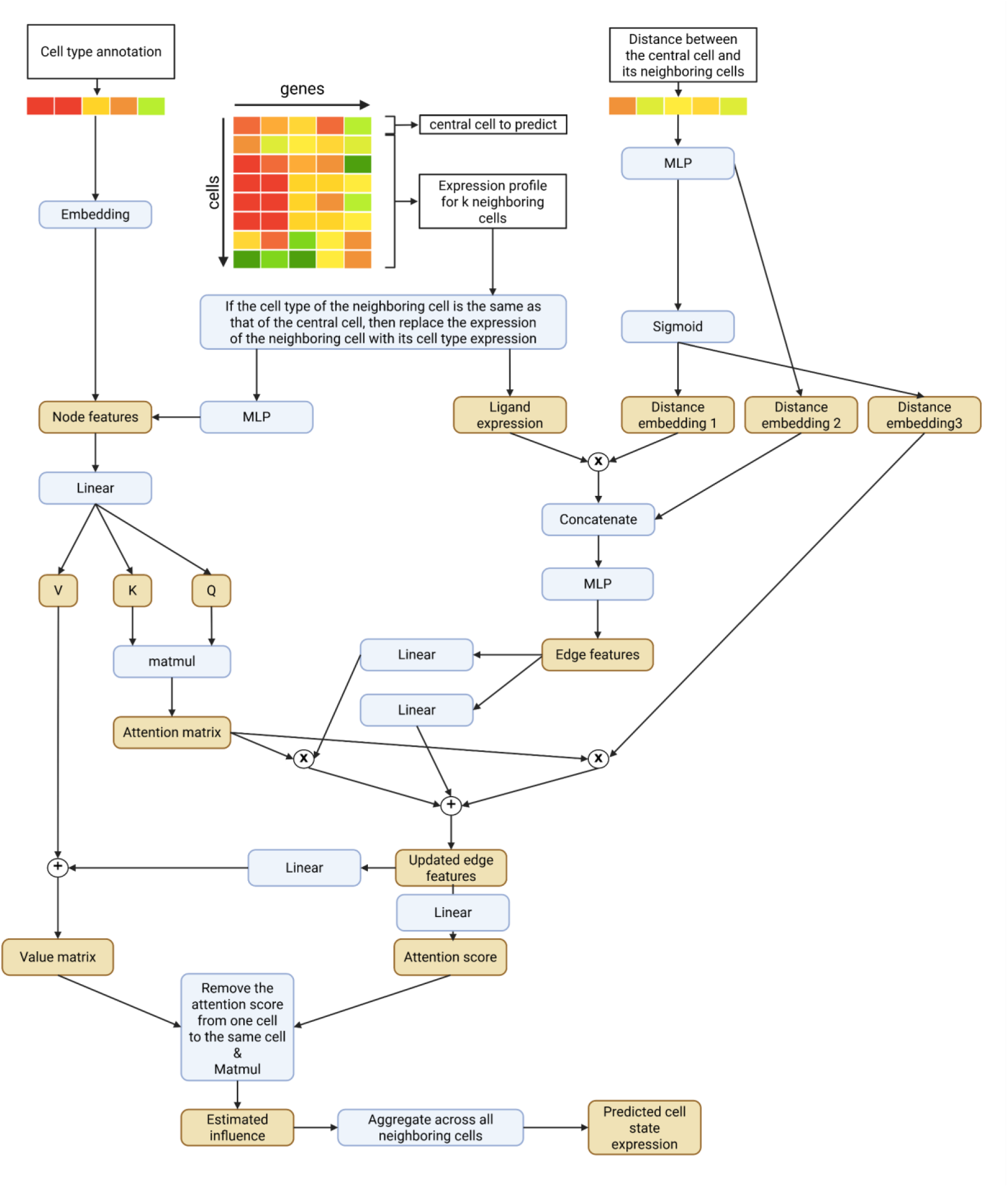
**Graphical illustration of the data processing workflow in GITIII.**

**Supplementary Fig. 2.**
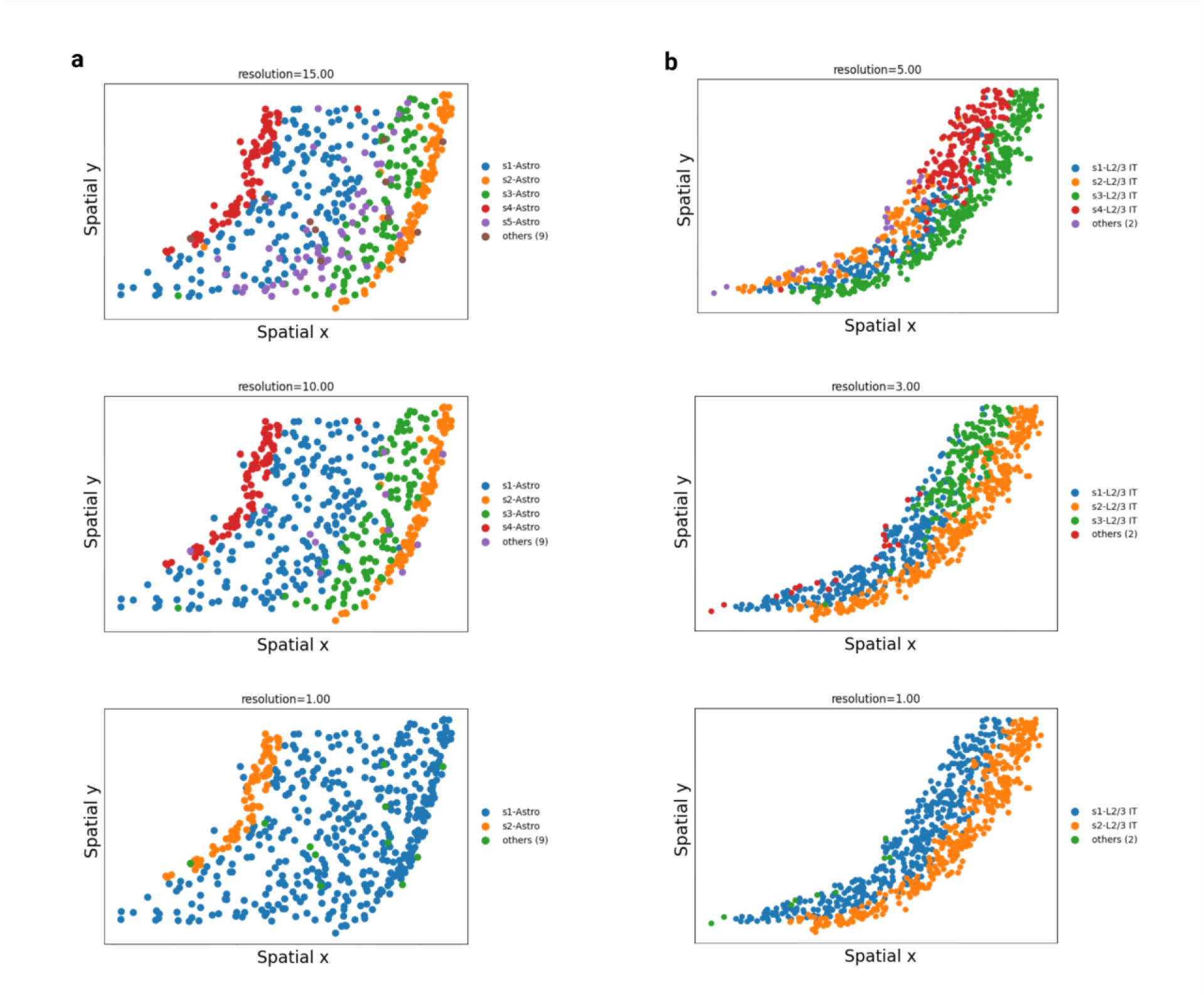
Spatial clustering by BANKSY in the mouse PMC dataset. **a,** Spatial distribution of astrocyte subgroups at different resolutions. Cells labeled as “others (x)” belong to small subgroups containing fewer than 20 astrocytes, where x denotes the number of such subgroups. **b,** Spatial distribution of L2/3 IT subgroups at different resolutions. Cells labeled as “others (x)” belong to small subgroups containing fewer than 20 L2/3 IT neurons, where x denotes the number of such subgroups.

**Supplementary Fig. 3.**
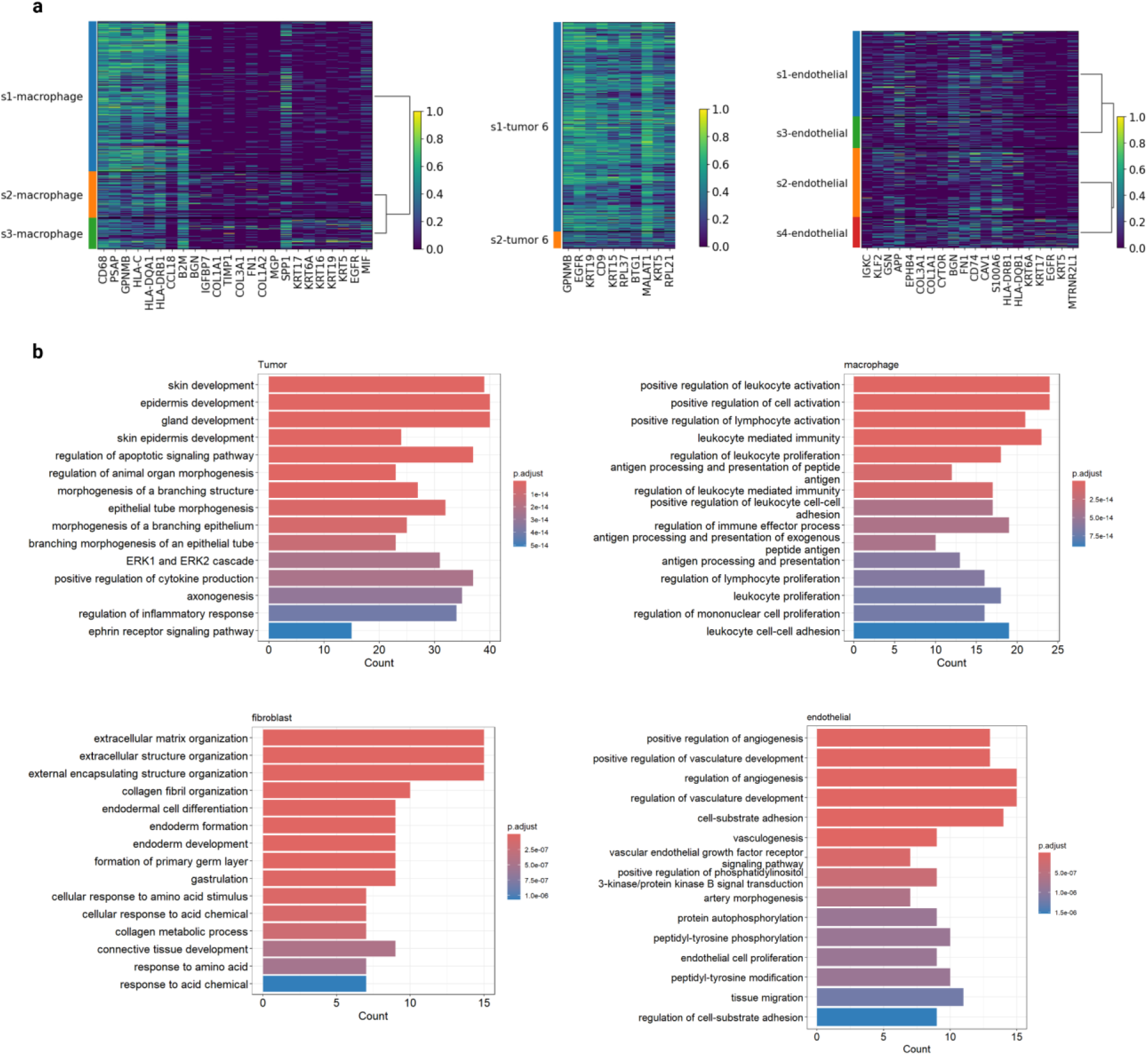
Heatmap of differential expressed genes and pathways of cell-type markers in the NSCLC dataset. **a,** Heatmap of the top differentially expressed genes in macrophage, endothelial, and tumor cell subgroups. **b,** GO enrichment analysis of the cell-type markers of tumor cell, macrophage, fibroblast, and endothelial cell.

**Supplementary Fig. 4.**
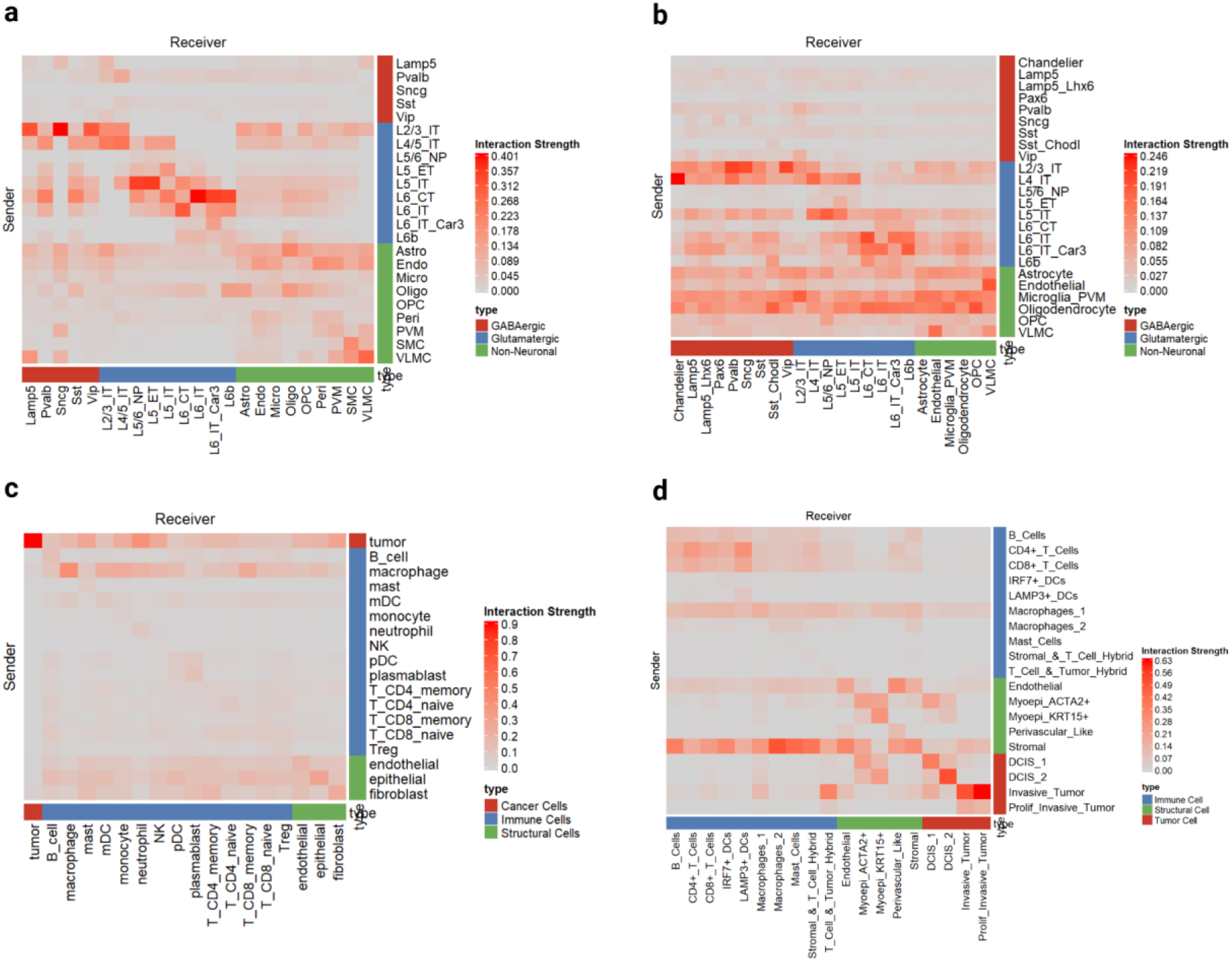
**Composition of sender cell types influencing each receiver cell type in the (a) mouse PMC, (b) SEA-AD, (c) NSCLC, and (d) breast cancer datasets.**

**Supplementary Fig. 5.**
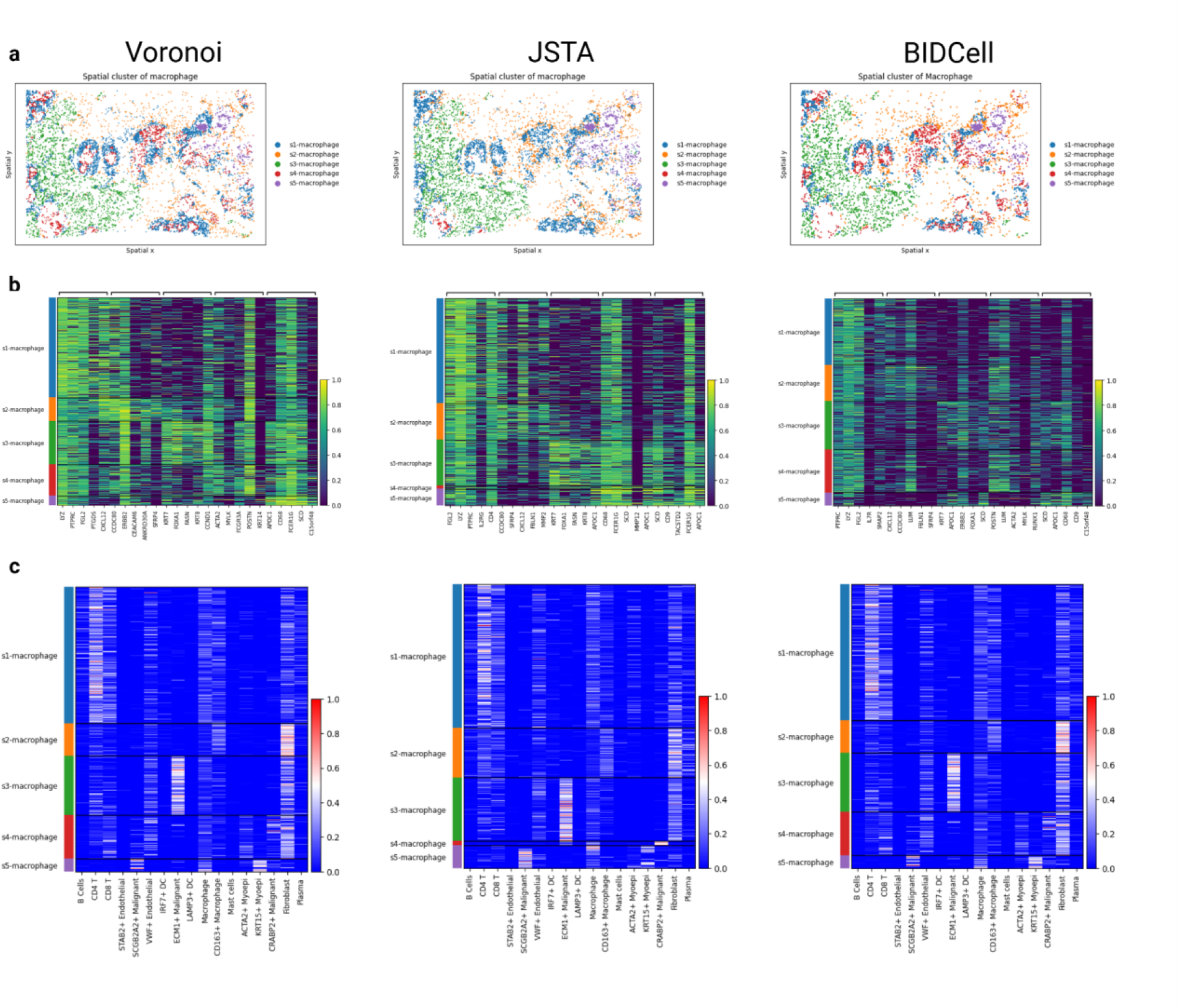
Comparison of clustering results of macrophages across data processed by different cell segmentation methods. **a,** Spatial distribution of subgroups. **b,** Heatmap of the top differentially expressed genes in subgroups. **c,** Neighboring cell type influence on subgroups, across three cell segmentation methods: Voronoi (left), JSTA (middle), and BIDCell (right).

**Supplementary Fig. 6.**
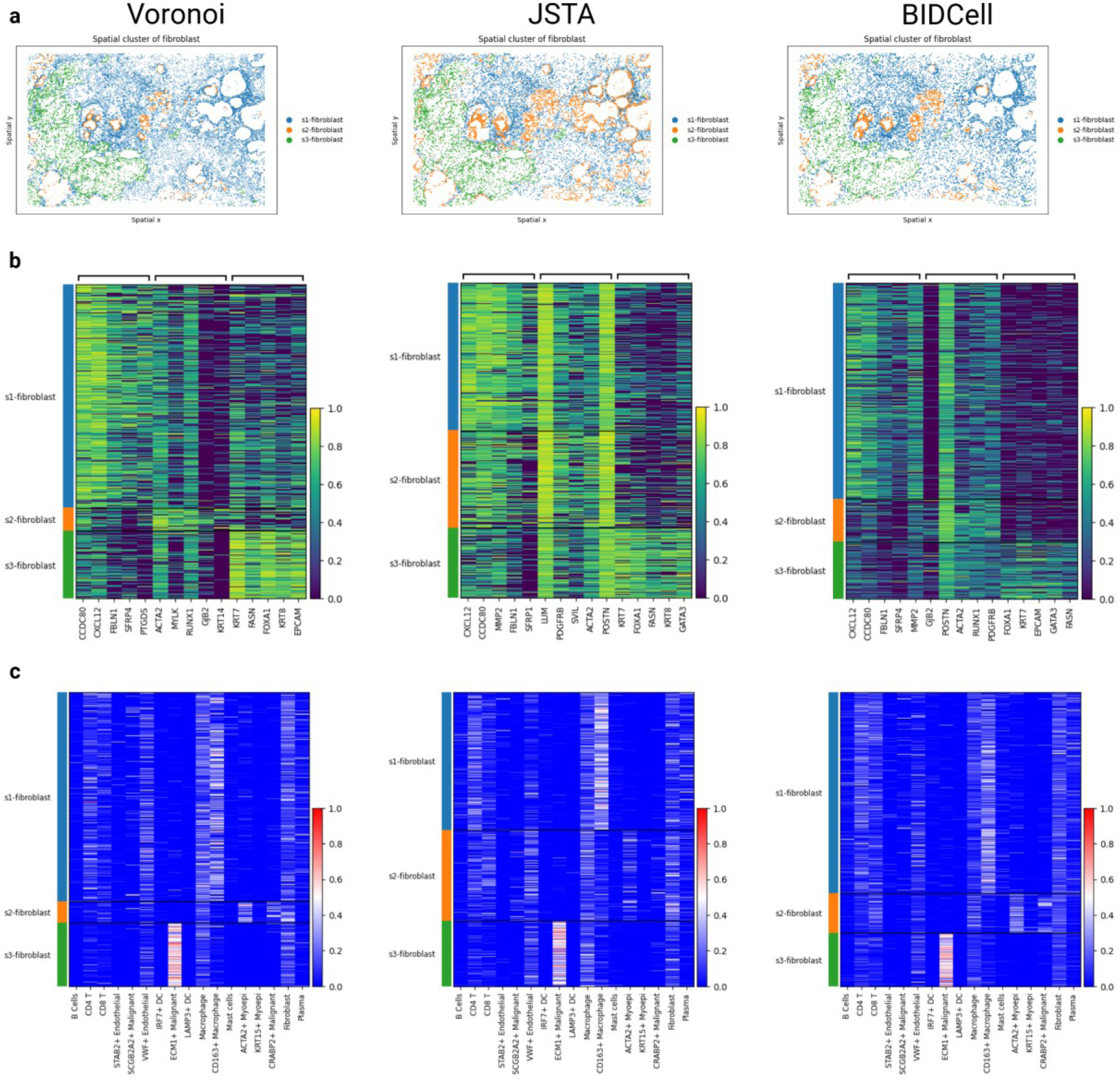
Comparison of clustering results of fibroblasts across data processed by different cell segmentation methods. **a,** Spatial distribution of subgroups. **b,** Heatmap of the top differentially expressed genes in subgroups. **c,** Neighboring cell type influence on subgroups, across three cell segmentation methods: Voronoi (left), JSTA (middle), and BIDCell (right).

**Supplementary Fig. 7.**
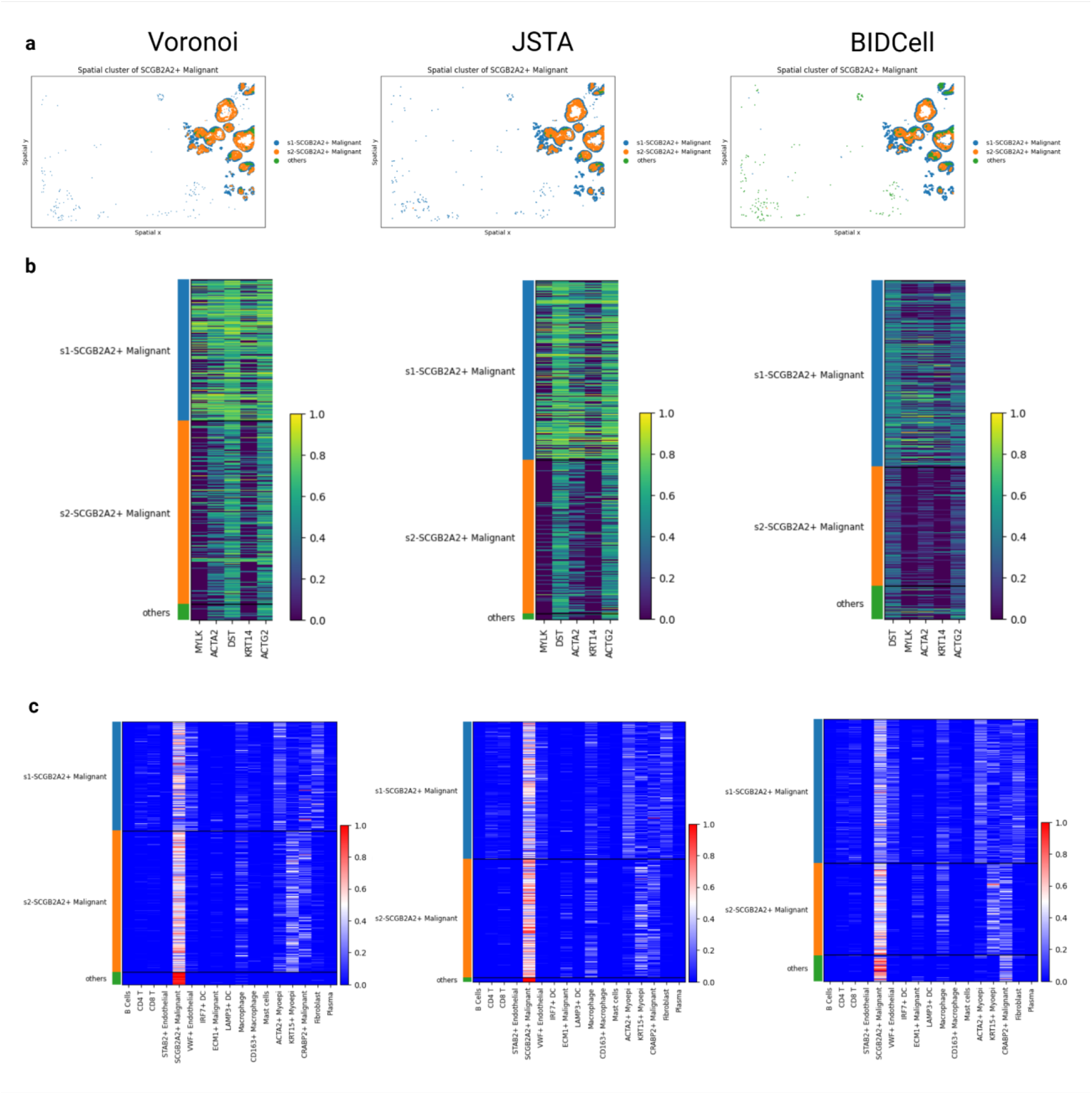
Comparison of clustering results of SCGB2A2+ malignant cells across data processed by different cell segmentation methods. **a,** Spatial distribution of subgroups. **b,** Heatmap of the top differentially expressed genes in subgroups. **c,** Neighboring cell type influence on subgroups, across three cell segmentation methods: Voronoi (left), JSTA (middle), and BIDCell (right).

**Supplementary Fig. 8.**
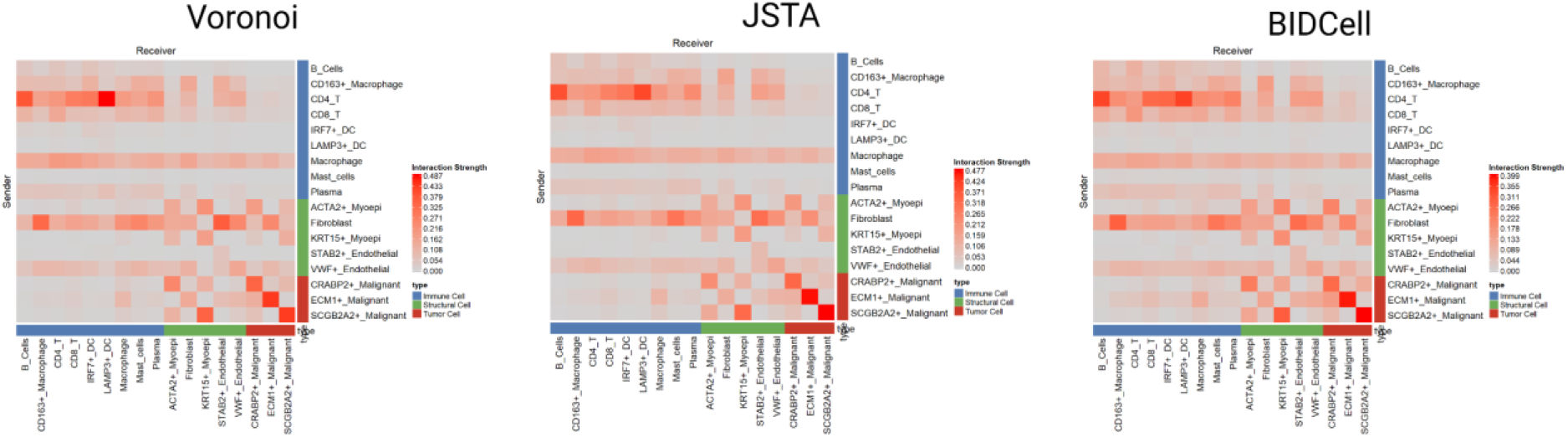
**Composition of sender cell types influencing each receiver cell type in the breast cancer data processed by three cell segmentation methods: Voronoi (left), JSTA (middle), and BIDCell (right)**

