## Supplementary material for "Inferring spatial single-cell-level interactions through interpreting cell state and niche correlations learned by self-supervised graph transformer": https://drive.google.com/file/d/1SJgG6oOVC1OgxTLeYypxDwqpb7-3LzAJ/view?usp=sharing

### 1 List of notations

#### 1.1 Model input

- The expression matrix $\text{Exp} \in \mathbb{R}^{k \times c}$, where $k$ represents the number of neighboring cells in the neighborhood, including the central cell, and $c$ is the number of measured genes in this dataset.
- The distance matrix distance $\in \mathbb{R}^{k \times k}$, describing the pairwise distances between the cells.
- The cell type vector $T \in \mathbb{R}^{k}$, where each entry corresponds to one of the $t$ cell types identified in this dataset.

#### 1.2 Embedding module

- The function of multi-layer perceptron (MLP), suppose the input is a tensor $x \in \mathbb{R}^{* \times c_1}$, where $*$ can be any value or sequence of values, $c_1$ is its number of features. And we want the output $\in \mathbb{R}^{* \times c_2}$, where $c_2$ is the number of dimensions we want to project $x$ on. The MLP uses two linear transformation with the GELU activation function to make the encoding. Specifically, output $= \text{GELU}(xW_1)W_2$, where $W_1$ and $W_2$ are the weight matrix of the two linear transformation.
- Node features are represented by the matrix $X \in \mathbb{R}^{k \times D_V}$, where k corresponds to the number of cells in a neighborhood, and $D_V$ denotes the embedding dimension of the nodes. Analogous to the input tensor of a language model, each cell in this context is analogous to a word (token), and the entire neighborhood of k cells is treated as a sentence comprising k words.
- Edge features are represented by the tensor $E \in \mathbb{R}^{k \times k \times D_E}$, where $D_E$ represents the embedding dimension of the edges. This tensor characterizes the interactions between every pair of cells within the k-cell neighborhood.
- Distance embeddings are represented by the tensor $A \in \mathbb{R}^{k \times k \times D_E}$, detailing the influence of the spatial distance between each pair of cells in the neighborhood.
- The function of mask in the node embedding section:

$$M(\mathbf{x}_j, c_j, c_i) = \begin{cases} \bar{\mathbf{x}}_{c_j}, & \text{if } c_j = c_i \\ \mathbf{x}_j, & \text{otherwise} \end{cases}$$

where:

- $\mathbf{x}_j$ is the gene expression profile of the neighboring cell $j$.
- $c_j$ is the cell type of the neighboring cell $j$.
- $c_i$ is the cell type of the central cell $i$.
- $\bar{\mathbf{x}}_{c_j}$ represents the average expression profile of all cells of type $c_j$.

#### 1.3 Downstream analysis

##### 1.3.1 Processing of the influence tensor

- Influence tensor $I \in \mathbb{R}^{N\times(k-1)\times c}$, where $N$ is the number of cells in the analyzed slide (tissue section), $c$ is the number of measured genes in the dataset and k-1 is the number of cells in the cell neighborhood used for prediction. This tensor characterizes how each cell's each gene's state is influenced by its top k-1 nearest neighbor cells. We can select the top 5 items along the second dimension for the UMAP visualization of CCI pairs (How one sender cell influences each gene's state in the corresponding receiver cell that forms an interaction cell pair).

- Proportionally normalized influence tensor $I_p \in \mathbb{R}^{N\times(k-1)\times c}$, which is the absolute value of the influence tensor $I$ with all items within the same second dimension sum up to 1. $I_p$ represents the proportional influence from the top k-1 nearest neighbors targeting each cell's each target gene.

- Proportionally normalized influence tensor averaged across all genes $I_s \in \mathbb{R}^{N\times(k-1)}$, which is the absolute value of the influence tensor $I$ with all items within the same second dimension sum up to 1. $I_s$ quantifies how each cell is proportionally influenced by its top k-1 nearest neighbors averaged across all target genes.

- Aggregated influence tensor $\mathbb{I} \in \mathbb{R}^{N\times t\times c}$, where $N$ refers to the number of cells in the analyzed slide, $t$ refers to the number of unique cell types in this slide, and $c$ donates the number of measured genes. This tensor characterizes how each cell's each gene's state is influenced by other cell types.

### 2 Distance embedding

#### 2.1 Technical details

Given a distance matrix distance that characterize the distance between each cell in the input cell neighborhood, we can calculate the distance feature tensor by:

$$\text{Distance\_features} = \text{stack}\left(\frac{1}{1+\text{distance}}, \frac{1}{1+\text{distance}^2}, \frac{1}{1+\sqrt{\text{distance}}}, e^{-\text{distance}}, e^{-\text{distance}^2}\right). \tag{1}$$

where Distance_features $\in \mathbb{R}^{k \times k \times 5}$ and "stack" means unsqueeze the last dimension for all the tensors and concatenate them along that dimension.

Then, the three distance embedding can be calculated by:

$$\text{DE1} = \text{MLP1}(\text{Distance\_features}) \quad (2)$$

$$\text{DE2} = \text{Sigmoid}(\text{MLP2}(\text{Distance\_features})) \quad (3)$$

$$\text{DE3} = \text{Sigmoid}(\text{MLP3}(\text{Distance\_features})) \quad (4)$$

$$\text{Sigmoid}(x) = \frac{1}{1 + e^{-x}} \quad (5)$$

where MLP1 and MLP3 transform the last dimension of features from 5 to the $D_E$, and MLP2 tranforms the last dimension of features from 5 to the number of measured ligands in the dataset #L.

The intuition of the above distance embedding calculation is described below: DE1 is a direct distance embedding that would be treated as the input of other neural networks to model complex non-linear distance's effect on CCI. DE2 and DE3 are designed to scale the ligand expression and the KQ attention tensor in the attention calculation, since we know that, in physics, the effect of distance on the strength of one LR signaling pathway is scaling. As a scaler, we applied a sigmoid function to them to make their values range from 0 to 1.

#### 2.2 Rationale on the use of distance information

The distance features we construct differ from the commonly used exponential decay diffusion model to better describe dynamic processes in biological systems. First, in cases where L-R pairs function only when cells are in direct contact, Fick's second law of diffusion is not applicable, as there is no ligand diffusion. Second, the diffusion equation assumes that ligand concentrations in the space are initially zero everywhere except at the source and that the space is infinite. These assumptions are often not met in biological systems. For example, tumor microenvironments may have a high local concentration of growth factors secreted by cancer cells. Additionally, in the blood-brain barrier, diffusion is constrained not only by the selective permeability of the brain but also by the limited spaces within the brain's structural compartments. Third, the diffusion equation assumes that no reactive processes affect the particles during diffusion. However, in tissues such as the brain, neurotransmitters crossing synaptic gaps are rapidly broken down by enzymes like acetylcholinesterase, significantly reducing their concentration and effectiveness. Fourth, The factors in the principles of ligand diffusion are not the same and unknown for different ligands. In summary, it is almost impossible to accurately estimate the distance decay and it is improper to use a fixed distance decay function for all LR pathways.

In addition, of note that the visualized distance decay trend in Fig. 2-5b is an average of all estimated influences, where the interacting strength within each distance interval is averaged in the plot. Though the interaction strength decays with distance on the general trend, it does not necessarily mean that interaction between closer cells must be stronger than cells with larger distances, or only direct contact CCI is ranking on the top. To illustrate, we plotted the the range of interaction strength targeting the RORB gene on the 'H20.33.001.CX28.MTG.02.007.1.02.03' slide in the AD MERFISH dataset (Fig. S4). We can observe that the situation exists where cells at further distances have stronger interaction strength than the cells at closer distances, which means that GITIII not only infers CCI according to the spatial organization.

Biologically, this makes sense because if there is little to no ligand-receptor expression targeting this gene, the interaction strength will remain zero or very weak, even if the two cells are in close proximity. However, the general trend shows that closer cells tend to exhibit stronger interaction strength. This is consistent with the physical fact that ligand concentration decreases exponentially with increasing distance due to the diffusion effect.

### 3 Edge embedding

The edge embeddings takes in 3 components: ligand expressions, DE1 and DE2. And it contains 3 steps: calculation of ligand level, ligand transformation, and integration with DE1.

Firstly, for different ligands, we compute the corresponding ligand level using the measured expressions of gene(s) that make up this ligand. Similar to Neuronchat, AND logic (i.e., geometric mean) is applied among different groups of genes that form one ligand; since the genes within the same group to produce one ligand are redundant for the same function, the OR logic (i.e., arithmetic mean) is applied.

The computed ligand level, denoted as ligand_level, initially has dimensions $\mathbb{R}^{k \times \#L}$. It is then expanded along the second dimension to the dimensions of $\mathbb{R}^{k \times k \times \#L}$, whereby each element along this second dimension replicates the original values. Subsequently, it is encoded by a linear transformation by:

$$\text{Ligand\_score} = \text{Linear}(\text{Ligand\_level} \times \text{DE2}). \tag{6}$$

where Linear donates a linear transformation from the dimension of $\#L$ to $D_E$.

Then, the edge features is crafted by integrating the ligand score with another transformation of the distance features DE1:

$$E = W(\text{concatenate}(\text{Ligand\_score,DE1})) \times \sqrt{D_E} \tag{7}$$

where W $\in \mathbb{R}^{2D_E \times D_E}$ represents a weight matrix that transforms the concatenated tensor with shape $\mathbb{R}^{k \times k \times 2D_E}$ to $\mathbb{R}^{k \times k \times D_E}$. The purpose of multiplying by $\sqrt{D_E}$ is to appropriately scale the edge features, thereby ensuring their comparability with the attention tensor that is subsequently computed within the encoder.

### 4 Graph attention calculation

#### 4.1 Preliminaries on tensor operation

To help the reader better understand the tensor operations including squeeze, reshape, and permute, here we provided a systematic illustration of these operations:

- **Permute** reorders the axes of a tensor according to a specified permutation, effectively changing the orientation of the data. Formally, for a permutation $\sigma$ of $\{1, 2, \ldots, n\}$, the permuted tensor $T'$ is defined as:

$$T'_{i_{\sigma(1)}, i_{\sigma(2)}, \ldots, i_{\sigma(n)}} = T_{i_1, i_2, \ldots, i_n}.$$

  For example, given a tensor $T \in \mathbb{R}^{2\times3}$ as

$$T = \begin{bmatrix} 1 & 2 & 3 \\ 4 & 5 & 6 \end{bmatrix},$$

  applying a permutation of axes with `permute(1, 0)` yields $T' \in \mathbb{R}^{3\times2}$:

$$T' = \begin{bmatrix} 1 & 4 \\ 2 & 5 \\ 3 & 6 \end{bmatrix}.$$

- **Reshape** changes the shape of a tensor without modifying the order or number of elements. The operation preserves data continuity and is valid as long as the total number of elements remains constant. Continuing from the example above, reshaping $T \in \mathbb{R}^{2\times3}$ into $T' \in \mathbb{R}^{3\times2}$ results in:

$$T' = \begin{bmatrix} 1 & 2 \\ 3 & 4 \\ 5 & 6 \end{bmatrix}.$$

- **Squeeze** removes singleton dimensions—those of size 1—thereby reducing the tensor's rank and simplifying its structure. This is particularly useful for eliminating redundant axes introduced during data preparation or broadcasting. For instance, a tensor $T \in \mathbb{R}^{1\times3\times1}$ given by

$$T = \begin{bmatrix} \begin{bmatrix} 1 \\ 2 \\ 3 \end{bmatrix} \end{bmatrix}$$

  becomes $T' \in \mathbb{R}^3$ after applying `squeeze()`:

$$T' = \begin{bmatrix} 1 & 2 & 3 \end{bmatrix}.$$

#### 4.2 Algorithm

##### 4.2.1 Calculation of the attention tensor

We first calculate the attention tensor generated from the node features:

$$K = (XW_K).\text{reshape}(k, D_A, 2D_E).\text{permute(2,0,1)} \quad (8)$$
$$Q = (XW_Q).\text{reshape}(k, D_A, 2D_E).\text{permute(2,1,0)} \quad (9)$$
$$\text{KQ} = (QK).\text{permute(1,2,0)} \quad (10)$$

Here, node features $X$ from the node embedding module are transformed into key (K), query (Q), and value (V), each has the shape of $\mathbb{R}^{k\times(2D_E\times D_A)}$, via linear transformations using learnable weight matrices $W_K, W_Q \in \mathbb{R}^{C\times(2D_E\times D_A)}$. Here, $D_A$ donates the dimension required for one-head attention computation, which is 8 as default. These matrices are then reshaped and permuted to create a multi-head attention tensor $\mathrm{KQ} \in \mathbb{R}^{2D_E\times k\times k}$, akin to the approach in the original attention mechanism paper. The attention tensor is permuted to align with the shape of the edge features $\in \mathbb{R}^{k\times k\times D_E}$.

This operation is designed to encapsulate latent relational information that is not directly measured in the dataset but is potentially inferable. For example, suppose gene B produces a ligand that is not captured in a single-cell-resolution spatial transcriptomics dataset. If its expression is strongly correlated with a measured gene A, or can be inferred from the set of measured genes, then gene B's information may be partially embedded within the variable $K$. Similarly, if gene D is an unmeasured receptor whose expression can be inferred from the measured gene set, then its information may be captured by the variable $Q$. After computing the attention tensor, the product $KQ$ can be used to infer the missing ligand–receptor (LR) signaling pathway between gene B and gene D. In summary, $KQ$ serves as a compensator for the loss of unmeasured LR gene information.

##### 4.2.2 Update edge features

Once the attention tensor is derived from the node features, we update the edge features through a composite process. This update integrates the newly calculated attention tensor with previous edge features and the distance embedding. This dynamic update procedure refines the edge features, capturing the complexities of cell-cell interaction (CCI), which is influenced by multiple factors. These include spatial proximity, both observed and unobserved ligand-receptor (LR) pairs, and additional elements such as the extracellular matrix, potentially governed by specific genes. This can be mathematically represented as follows:

$$E = \mathrm{gelu}(\rho(\mathrm{KQ}[:,:,:D_E/2] \odot EW_{Ew}) + EW_{Eb} + \mathrm{KQ}[:,:,D_E/2:] \odot \mathrm{DE3}) \tag{11}$$

$$\rho(x) = \sqrt{\mathrm{relu}(x)} - \sqrt{\mathrm{relu}(-x)} \tag{12}$$

where $\odot$ denotes element-wise multiplication of tensors using the Hadamard product, $[:,:,:D_E/2]$ donates slicing the tensor along the last dimension, $\rho$ represents the signed-square-root function, which stabilizes training by moderating the impact of large input values, and relu refers to the Rectified Linear Activation function. Learnable weight matrices $W_{Ew}$ and $W_{Eb}$ are dimensioned in $\mathbb{R}^{D_E\times D_E}$, and after updating, the edge features $E$ reside in $\mathbb{R}^{k\times k\times D_E}$.

Specifically, the term $\mathrm{KQ}[:,:,:D_E/2] \odot EW_{Ew}$ captures the impact of ligands on the target cells, where the ECM and receptor information extracted from the node features ($\mathrm{KQ}[:,:,:D_E/2]$) is multiplied by the ligands' representation $EW_{Ew}$. In addition, $EW_{Eb}$ acts as an additional encoding layer for ligand information. Furthermore, $\mathrm{KQ}[:,:,D_E/2:] \odot \mathrm{DE}$ represents the LR information that might not be explicitly measured in the dataset but can be inferred from other measured genes in the node features.

The intuition of the above attention score calculation is described below:

- The D/2 operation simply splits the large KQ attention tensor that contains the receiver cell's information, the sender cell's information, and the unmeasured LR's information, into two parts.

- The term $\mathrm{KQ}[:,:,:D_E/2] \odot EW_{Ew}$ captures the impact of measured ligands on the target cells. A biological intuition is that, the edge feature $E$ that contains the information of measured ligand and distance information, after being transformed by a weight matrix, is adjusted by $\mathrm{KQ}[:,:,:D_E/2]$ that contains the receptor information on the receiver cell. So this term represents the effect of known ligand by integrating the known ligand, receiver cell, and distance information.

- $\mathrm{KQ}[:,:,D_E/2:] \odot \mathrm{DE3}$ is designed to capture the unmeasured LR signaling pathways' influence on the receiver cell, where the $\mathrm{KQ}[:,:,D_E/2:]$ that contains unmeasured LR information is scaled by the distance embedding DE3.

- The $EW_{Eb}$ term can be viewed as a tranformation of the original edge features.

- Together, these two parts use neural networks to estimate the cell-cell interactions mediated by the measured and the unmeasured ligand-receptor interactions, as well as distance's effect.

###### 4.2.3 Influence tensor calculation

Then, the attention score $\alpha \in \mathbb{R}^{k\times k}$ is computed by:

$$\alpha = \mathrm{softmax}((EW_A).\mathrm{squeeze}(\mathrm{dim}\text{=-1}), \mathrm{dim}\text{=-1}) \tag{13}$$

where $W_A \in \mathbb{R}^{D_E\times 1}$, and "squeeze" refers to removing the last dimension with length 1. After that, the softmax function is applied to normalize the attention score for each of the cells in the cell neighborhood (along the -1 dimension).

After that, the influences of the 1nd to the k-th tokens (neighboring cells) on the central cell (0-th token) from node features $i_n$ and edge features $i_e$ are calculated by the following equations:

$$\alpha' = \alpha[0, 1:] \tag{14}$$

$$i_n = \alpha\mathbf{1'} \bigodot X\,[1:,:]\,W_V \tag{15}$$

$$i_e = \alpha' E[0, 1:,:]W_{En} \tag{16}$$

Of note that the subscript starts from 0, which is the form in the code.

Initially, the attention scores $\alpha$ are truncated to produce a vector $\alpha' \in \mathbb{R}^{1\times(k-1)}$ that represent the impact of all neighboring cells on the central cell to predict. Subsequently, the node and edge features associated with these cells are extracted and transformed using the weight matrices $W_V \in \mathbb{R}^{C\times C}$ and $W_{En} \in \mathbb{R}^{D_E\times C}$, respectively.

Finally, the contributions from both node and edge features are combined and processed through an MLP head to yield the influence matrix $i \in \mathbb{R}^{(k-1)\times c}$. The matrix $i$ is then summed along the first dimension to derive the final prediction of the cell state $\hat{y} \in \mathbb{R}^c$, which is then compared to the measured cell state expression $y \in \mathbb{R}^c$ by mean square error to update model parameter:

$$i = \text{MLP}(\text{concatenate}(i_n, i_e)) \tag{17}$$
$$\hat{y} = \text{sum}(i, \text{dim} = 0) \tag{18}$$

Concatenating all influence matrix for all cells within one slide results in the influence tensor $I \in \mathbb{R}^{N \times (k-1) \times c}$, where $N$ indicates the number of cells within a slide.

The whole pipeline, from input data to the estimated influence, is demonstrated in the graphical illustrations Supplementary Fig. 1.

### 5 Justification for using the influence tensor as a measure of CCI influence

In a graph attention network (GAT), each layer $l$ computes:

$$\mathbf{h}_i^{(l+1)} = \sigma\left(\sum_{j \in \mathcal{N}(i)} \alpha_{ij}^{(l)} \mathbf{W}^{(l)} \mathbf{h}_j^{(l)}\right), \tag{19}$$

where

- $\mathbf{h}_i^{(l)}$ is the hidden representation of node $i$ at layer $l$,
- $\alpha_{ij}^{(l)}$ is the attention score for edge $(j, i)$,
- $\mathbf{W}^{(l)}$ is a learnable linear transformation,
- $\sigma(\cdot)$ is a nonlinear activation (e.g., ReLU).

To simplify, we assume that there is only one attention head. After $L$ layers, GAT includes a final linear transformation $\mathbf{W}_{\text{out}}$ (and possibly another activation $\phi$):

$$\mathbf{z}_i = \mathbf{W}_{\text{out}} \mathbf{h}_i^{(L)} \quad \text{or} \quad \mathbf{z}_i = \phi\Big(\mathbf{W}_{\text{out}} \mathbf{h}_i^{(L)}\Big), \tag{20}$$

where $\mathbf{h}_i^{(L)}$ is the hidden state of node $i$ at the last GAT layer $L$. Thus, the final output $\mathbf{z}_i$ in a multi-layer GAT depends on:

1. The attention scores $\alpha_{ij}^{(l)}$ at each layer $l$ from 0 to $L$,
2. The hidden states $\mathbf{h}_j^{(l)}$, which themselves are outputs of previous layers,
3. Nonlinear activations $\sigma$,
4. The final matrix $\mathbf{W}_{\text{out}}$.

#### 5.1 Information aggregation

The neighborhood aggregation is:

$$\sum_{j\in\mathcal{N}(i)} \alpha_{ij}^{(l)}\mathbf{W}^{(l)}\mathbf{h}_j^{(l)}, \tag{21}$$

which then passes through the nonlinear activation $\sigma$. Consequently, the final hidden state $\mathbf{h}_i^{(l+1)}$ is not directly proportional to $\alpha_{ij}^{(l)}$; it is a nonlinear function of both $\alpha_{ij}^{(l)}$ and the neighbor embeddings $\mathbf{h}_j^{(l)}$.

In other words, in attention-based GNN models, the neighborhood information is aggregated to the central node by summing the multiplication of attention scores and values over all neighboring nodes, so the multiplication of attention scores and values is a better indicator of the influence of other nodes (cells) compared to solely attention scores.

#### 5.2 Multiple layers compound the complexity.

Traditional GNN models contain multiple GNN layers, the influence from the neighboring nodes to the central node is calculated multiple times in each GNN layer, making it hard to explicitly obtain the influence from other cells to the central receiver cell. So in GITIII, we adopted a single-layer GNN model to obtain the influence from neighboring sender cells instead of using multiple layers.

#### 5.3 Final outputs include a further transformation by $\mathbf{W}_{\text{out}}$ and $\sigma$.

Since

$$\mathbf{z}_i = \mathbf{W}_{\text{out}}\sigma\left(\sum_{j\in\mathcal{N}(i)} \alpha_{ij}^{(L-1)}\mathbf{W}^{(L-1)}\mathbf{h}_j^{(L-1)}\right). \tag{22}$$

Even the last-layer attention $\alpha_{ij}^{(L-1)}$ is subsequently transformed by both the nonlinear activation $\sigma$ and the additional matrix $\mathbf{W}_{\text{out}}$. Therefore, the final output $\mathbf{z}_i$ is not simply a linear combination of $\alpha_{ij}^{(L-1)}\mathbf{h}_j^{(L-1)}$. The presence of $\sigma$ and $\mathbf{W}_{\text{out}}$ further warps this relationship, making attention alone insufficient to interpret each neighbor's exact influence.

In other words, attention-based GNN models apply a non-linear transformation after aggregating the neighboring node information using attention mechanisms, which means the influence from neighboring cells is not the multiplication of attention scores and values due to non-linearity. For GCN-based models, the interpretation would also be better if there is no final non-linear transformation.

So we simply removed the non-linear transformation to make sure that, in theory, the multiplication of attention scores and values in our single-layer, FFN-free GNN encoder represents the CCI influence of neighboring cells to the central receiver cell.

#### 5.4 Our approach

GITIII only uses a single attention-based layer without additional transformations, the model output for node $i$ can be as simple as:

$$\mathbf{z}_i = \sum_{j \in \mathcal{N}(i)} \alpha_{ij} \mathbf{v}_j, \tag{23}$$

where $\mathbf{v}_j$ is the "value" vector for node $j$. In this scenario, the contribution of each neighbor $j$ is literally $\alpha_{ij}\mathbf{v}_j$. Because there is no further activation or matrix multiplication, attention $\times$ value in this single-layer setup directly quantifies neighbor $j$'s influence on node $i$'s final output.

Hence, in our single-layer direct-sum approach, the neighbor's contribution is indeed "attention $\times$ value." By contrast, in a multi-layer GAT with final transformations, attention scores alone are insufficient to explain the final output because of the intervening nonlinearities and final $\mathbf{W}_{\text{out}}$.

In GNN models, multiple layers are typically used, with each layer repeatedly propagating information from neighboring nodes to a central node. This multi-layer propagation makes it challenging to explicitly quantify the influence of individual neighboring cells on a central receiver cell. To address this, GITIII employs a single-layer GNN architecture to directly calculate the influence from neighboring sender cells to the central receiver cell. In the attention-based GNN models, neighborhood information is aggregated as a weighted sum, where each term is the product of the attention score and the corresponding value of neighboring node. Without applying additional nonlinear transformations, this product directly represents the individual contribution of each neighboring sender cell. Therefore, we did not apply any nonlinear transformation after aggregation to preserve the interpretability of the influence.

### 6 Enrichment analysis of cell-type-specific genes: evidence against artefactual lateral spillover

We performed enrichment analyses on cell-type specific genes for tumor cells, macrophages, fibroblasts, and endothelial cells. Compared to the pathways enriched for genes significantly influenced by CCI between tumor cells and the other three cell types, the cell-type-specific gene enriched pathways are largely different. For example, tumor-specific genes are enriched in pathways such as apoptotic signaling, ERK1 and ERK2 cascade that regulates a wide variety of cellular processes, including proliferation, differentiation, and survival, as well as inflammatory response (Supplementary Fig. 3b). In contrast, genes in endothelial cells influenced by tumor cells are enriched in pathways related to cytoskeleton and fiber growth (Fig. 4k), while genes in macrophages influenced by tumor cells are involved in apoptotic processes and extracellular organelle dynamics (Fig. 4l). These show little overlap with the tumor-specific gene pathways. Similarly, macrophage-specific genes are enriched for immunity-related pathways, including lymphocyte and leukocyte activation, and antigen processing and presentation (Supplementary Fig. 3b). In contrast, genes in tumor cells influenced by macrophages are enriched in pathways related to cell cycle, chromosome organization, and p53 signaling (Fig. 4j), again showing little to no overlap. Lastly, fibroblast-specific genes are enriched in extracellular matrix (ECM) organization and connective tissue development pathways (Supplementary Fig. 3b), while genes in tumor cells influenced by fibroblasts are enriched in ECM remodeling pathway (Fig. 4i). This is consistent with the literature showing that cancer-associated fibroblasts (CAFs) contribute to ECM modeling in tumor cells. Taken together, these results indicate that the pathways enriched in sender cell-type-specific genes are largely distinct from those enriched in CCI-influenced genes in the receiver cell types. This supports the conclusion that the CCI-related biological processes we found are unlikely to be artefacts of lateral spillover.

### 7 Benchmarking

**GAT.** Graph Attention Network (GAT) leverages the mechanism of attention in graph neural networks to focus on important neighbors. Like GCNs, GATs operate on graphs, but instead of treating all neighbors equally, they learn to weigh the importance of each neighbor's features dynamically. The key update rule in a GAT can be formulated as follows:

$$H^{(l)} = \sigma\left(\sum_{j\in\mathcal{N}_i} \alpha_{ij}^{(l)} W^{(l)} H_j^{(l-1)}\right) \tag{24}$$

where $H^{(l)}$ denotes the node representations at layer $l$, and $H_j^{(l-1)}$ is the representation of the $j$-th neighbor node from the previous layer. The coefficients $\alpha_{ij}^{(l)}$ are attention scores computed for each edge connecting node $i$ and its neighbor $j$ at layer $l$, determining the importance of node $j$'s features to node $i$. These attention scores are computed as:

$$\alpha_{ij}^{(l)} = \frac{\exp(\text{LeakyReLU}(\mathbf{a}^T[W^{(l)}H_i^{(l-1)} \| W^{(l)}H_j^{(l-1)}]))}{\sum_{k\in\mathcal{N}_i} \exp(\text{LeakyReLU}(\mathbf{a}^T[W^{(l)}H_k^{(l-1)} \| W^{(l)}H_j^{(l-1)}]))} \tag{25}$$

where $\mathbf{a}$ is a weight vector learned during the training process, $\|$ denotes concatenation, and the function $\sigma(\cdot)$ is LeakyReLU (cite here). Similarly, the dimensions and embeddings of node features and the optimizer are the same as in our model. We stacked two layers of GAT in our implementation. To build the spatial graph, we define an edge to exist between two cells if the Euclidean distance between the two cells is less than a neighborhood size. We set the neighborhood size to a set of values - [10, 30, 50, 75, 100, 150, 200, 250, 300, 400] pixels, and pick the one that explained the most intra-cell-type variance for benchmarking. This model does not identify statistically significant CCI networks between cell types, so it is only benchmarked through the criterion of the explained intra-cell-type variance. It can be applied across all 4 datasets.

**NCEM-GCN.** We set the node embedding dimension to 256 and construct the edge by setting the neighborhood size to a set of values - [10, 30, 50, 75, 100, 150, 200, 250, 300, 400] pixels, and pick the one that explained the most intra-cell-type variance for benchmarking. To make comparison and keep consistency, we set the loss function to the same as our model by removing the variance prediction term in the original paper. This model does not identify statistically significant CCI networks between cell types, so it is only benchmarked through the criterion of the explained intra-cell-type variance. It can be applied across all 4 datasets.

**COMMOT.** We run the COMMOT on the available datasets where measured ligand-receptor pairs are available in its built-in ligand-receptor database. The estimated received signal for each pathway for each cell is used to make a linear regression to predict its cell state expression. The prediction is then used for the calculation of the explained intra-cell-type variance. This model does not identify statistically significant CCI networks between cell types, so it is only benchmarked through the criterion of the explained intra-cell-type variance. It can be applied in all datasets excluding the AD dataset due to the lack of ligand-receptor pair for genes measured in the dataset.

**HoloNet (Multi-view Graph Convolutional Network).** In the implementation of the Multi-view Graph Convolutional Network (mv-GCN), we adopt the same embedding and model architecture as used in HoloNet. However, we omit the receptor expression information of the target cell in our predictions and use the ligand expression and the distance scaler purely derived from the chemical diffusion process as edge features. This exclusion is strategic in that it prevents the model from merely learning to decode from the product of ligand and receptor interactions to predict the expression of the cell to predict, as well as keep consistent with our model. Other methods like scaling the edge features by cosine similarity and edge features normalization were retained. The embedding dimension and the optimizer were the same as in our model. In addition, to make comparison and keep consistency, we only retain the loss function of cell state prediction. Furthermore, we assess a range of diffusion region diameters for a single cell—[100, 200, 255 (the default in many previous studies), 300, 400, 600, 1000, 5000, 10000, 50000, 100000] $\mu m$—and select the one that yields the lowest MSE loss on the validation set for benchmarking. In addition, to keep consistency, we modify the loss function of NCEM-GCN to the same as ours that reconstructs the cell state expressions instead of the measured expressions. This model does not identify statistically significant CCI networks between cell types, so it is only benchmarked through the criterion of the explained intra-cell-type variance. It can be applied in all datasets excluding the AD dataset due to the lack of ligand-receptor pair for genes measured in the dataset.

**Original Graph Transformer.** We apply the same model architecture as used in SiGra. To be more specific, the Graph Transformer employs a convolutional layer with a multihead attention mechanism as its core component. For a given node $v_i$, the propagation from layer $l$ to layer $l+1$ in a Graph Transformer is defined by the following equation:

$$h_i^{(l+1)} = \mathrm{ReLU}\left(W^{(l)}h_i^{(l)} + \sum_{v_j \in \mathcal{N}(v_i)} \alpha_{ij} V_j^{(l)}\right) \tag{26}$$

where ReLU is the rectified linear unit used as the nonlinear activation function. In this formulation, $h_i^{(l+1)}$ denotes the updated feature vector of node $v_i$ at layer $l+1$, and $h_i^{(l)}$ is the feature vector from the previous layer. The attention mechanism is crucial for weighing the influence of each neighbor $v_j$ in the neighborhood $\mathcal{N}(v_i)$. The attention coefficients $\alpha_{ij}$ are computed as follows:

$$\alpha_{ij} = \mathrm{softmax}\left(\frac{Q_i^{(l)}(K_j^{(l)})^T}{\sqrt{\dim(h_i^{(l)})}}\right) \tag{27}$$

where the query $Q_i^{(l)}$, key $K_j^{(l)}$, and value $V_j^{(l)}$ for the nodes are computed by linear transformation's weight matrix $W$ and $b$ as:

$$Q_i^{(l)} = W_Q^{(l)} h_i^{(l)} + b_Q^{(l)} \tag{28}$$

$$K_j^{(l)} = W_K^{(l)} h_j^{(l)} + b_K^{(l)} \tag{29}$$

$$V_j^{(l)} = W_V^{(l)} h_j^{(l)} + b_V^{(l)} \tag{30}$$

The attention scores are normalized using the softmax function, ensuring they sum to one. The Graph Transformer applies the multihead attention mechanism, where multiple sets of $Q$, $K$, and $V$ are learned and concatenated, allowing the model to attend to information from different representation subspaces at different positions in the graph simultaneously. In our implementation, the embedding dimension $h_i$, the methods to enumerate different distance thresholds and construct the sparse spatial graph and acquire the neighbors of each cell, the number of graph transformer layers, and the optimizer are the same as those used in our model. To build the spatial graph, we define an edge to exist between two cells if the Euclidean distance between the two cells is less than a neighborhood size. We set the neighborhood size to a set of values - [10, 30, 50, 75, 100, 150, 200, 250, 300, 400] pixels, and pick the one that explained the most intra-cell-type variance for benchmarking. This model does not identify statistically significant CCI networks between cell types, so it is only benchmarked through the criterion of the explained intra-cell-type variance. It can be applied across all 4 datasets.

**Models used in the cell-type level benchmarking.** We compare the aggregated CCI networks between cell types generated by GITIII, SpaTalk, COMMOT, stLearn, CellphoneDB (v5), CellChat (v2), SOApy, and Giotto, if applicable, using the metric described in Methods. For CellphoneDB and CellChat, their CCI analysis algorithms for spatial transcriptomics data were used.

For methods that aggregate CCI strength across different ligand–receptor (LR) gene pairs or varying numbers of cells—including `COMMOT`, `stLearn`, `SOApy`, and `CellChat`—we used their respective built-in functions to compute the overall CCI network strength between cell types. Specifically, in `COMMOT` we applied `ct.tl.cluster_communication` and accessed the results via `adata.uns['commot_cluster-subclass-user_database-total-total'][ 'communication_matrix']`. In `stLearn`, we extracted the values from `adata.uns[f"per_lr _cci_{cell_type}"]`. In `SOApy`, we extract the CCI network strength from the function `sp.pl.show_ccc_netplot(adata, lr_type='contact')` and `sp.pl.show_ccc_netplot( adata, lr_type='secretory')`. And in `CellChat`, we used the `aggregateNet` function. For methods that do not provide a built-in aggregation function, we computed the overall network strength by first select the significant results with adjusted p-value$<$0.05, then linearly scaling the interaction strength of each ligand–receptor (LR) pair such that the total strength across all cell type pairs for that LR pair sums to 1. After that, we averaged the corresponding CCI network strengths for each cell type pair across all LR pairs. To be more specific, for `SpaTalk`, we use the `obj@lrpair$score` from the output of the `dec_cci_all` function as the interaction strength; For `stLearn`, we use the `adata.uns["per_lr_cci_raw_cell_type"]` from the result of the function `stlearn.tl.cci.run_cci` as the interaction strength; For `Giotto`, we use the `PI` value from the results of the function `spatCellCellcom` as the interaction strength; For `CellphoneDB (v5)`, we use the `'interaction_scores'` from the results of the function `cpdb_statistical_analysis_method.call` as the interaction strength.

Due to the differences in the built-in ligand-receptor databases and the fact that some datasets do not have ligand-receptor pairs in their measured gene panels, SpaTalk is only applicable in the NSCLC dataset, while CellphoneDB and CellChat can be applied to both the NSCLC and BC datasets. In calculating the SCC between network strength and the distance between two cell types, we normalize the network strength by dividing it by the sum of all strength so that the total strength sum up to 1. In addition, the first two MERFISH datasets contain too few genes, and all methods and databases identified fewer than 10 ligand-receptor pair genes in the two datasets. It is too biased to use the CCI network between cell types aggregated from less than 10 LR pairs to represent the overall CCI that happens through over 2 thousand LR pairs. Besides, some methods are not applicable due to no ligand-receptor pair genes are detected. Due to the two reasons described above, we did not use these two datasets for cell-type-level benchmarking.

### 8 Evaluation of cell segmentation on the performance of GITIII

We assessed the robustness of GITIII to difference in cell segmentation in the breast cancer Xenium dataset. Specifically, we applied GITIII to the data processed by three cell segmentation methods: (1) classical Voronoi expansion, (2) JSTA, a joint cell segmentation and cell type annotation approach based on an extension of the watershed algorithm, and (3) BIDCell, a biologically-informed deep learning approach. Cell type annotations were assigned using scClassify, with detailed processing steps described previously. Because the number and labels of cell types were consistent across the three segmentation datasets, but differed from those in the original dataset, we focused our evaluation on the consistency of CCI results inferred by GITIII across the three datasets. In the first replicate slide from sample 1, Voronoi, JSTA and BIDCell produced 106,227, 107,131, and 103,209 cells, respectively.

At the single-cell level, we evaluated the robustness of GITIII by comparing the spatial organization, gene expression, and interacting cell types within the local microenvironment of CCI-informed subgroups for macrophages, fibroblasts (corresponding to stromal cells in Fig. 5d), and SCGB2A2+ malignant cells (corresponding to DCIS in Fig. 5d) across the three datasets. Macrophages were clustered into five subgroups with broadly consistent spatial distributions (Supplementary Fig. 5a), in agreement with the spatial patterns in Fig. 5e. The top differentially expressed genes in these subgroups showed largely overlap across datasets (Supplementary Fig. 5b). Subgroup 1 was mainly influenced by CD4 T cells, subgroup 2 by fibroblasts, and subgroup 3 by ECM1+ malignant cells (corresponding to invasive tumor in Fig. 5g; Supplementary Fig. 5c). Similarly, fibroblasts were consistently clustered into three subgroups with distinct expression profiles, with subgroup 3 mainly influenced by ECM1+ malignant cells in all datasets (Supplementary Fig. 6). For SCGB2A2+ malignant cells, we identified a group of cells located on the tumor boundary (subgroup 1) that consistently showed increased expression of *MYLK*, *ACTA2*, and *KRT14* (Supplementary Fig. 7). At the cell-type level, we compared the composition of interacting cell types for each receiver cell type across the three datasets and found that the normalized interaction strength was generally consistent for all cell-type pairs (Supplementary Fig. 8). Together, these results demonstrate that GITIII is robust to differences in cell segmentation.
